# Induction of synthetic apomixis in two sorghum hybrids enables seed yield and genotype preservation over multiple generations

**DOI:** 10.1101/2025.07.02.662806

**Authors:** Marissa K. Simon, Li Yuan, Ping Che, Kevin Day, Todd Jones, Ian D. Godwin, Anna M. G. Koltunow, Marc C. Albertsen

## Abstract

Apomixis, a process of clonal reproduction through seed, has the potential to significantly change agriculture by enabling a clonal seed propagation system for hybrid crops. Here, we demonstrate that hybrid seed from synthetically induced apomictic sorghum hybrids can be generated and maintained across multiple seed generations. This was achieved through the combination of avoidance of meiosis and induced parthenogenesis. Avoidance of meiosis was generated by the CRISPR/Cas9 knockout of the sorghum meiosis genes *Spo11*, *Rec8*, and *OsdL1* and *OsdL3*. Parthenogenesis was induced in the resultant diploid egg cell using a maize egg cell promoter to express the *Cenchrus ASGR-BBML2* gene coding sequence. Two strategies incorporating these components were used to induce synthetic apomixis in two different sorghum hybrids. Each hybrid used Tx623 as a female parent and either Tx430 or the African landrace Macia as a male parent. Seed yields in the induced apomictic hybrids were consistent and stable for multiple generations following self-pollination but reduced relative to the sexual hybrids. Sorghum contains two copies of the *Osd1* gene that function in meiotic non-reduction. CRISPR/Cas9 knockout of both *OsdL1* and *OsdL3* loci was sufficient to produce clonal hybrid progeny in conjunction with the other apomixis induction components, but this led to a significant reduction in seed set. By contrast, a single in-frame edit of either *OsdL1* or *OsdL3* significantly improved seed set of clonal hybrid progeny. Fine-tuning *OsdL* activity appears to be essential to optimizing fertility. As the efficiency of seed set in the induced synthetic sorghum apomicts was lower than that of the sexual hybrid control, additional improvements are required to unlock the agronomic potential of synthetically induced apomictic sorghum in the field.

## Introduction

Sexual reproduction in flowering plants leads to seed formation and is essential for the creation of genetic diversity through recombination during gametogenesis and subsequent parental gamete fusion during fertilization. However, some species can form seed asexually via apomixis where the maternal genotype is preserved in the progeny (Nogler, 1984). The developmental pathways resulting in apomictic seed vary in different apomictic species (Koltunow & Grossniklaus, 2003). Although apomixis occurs naturally in hundreds of plant species, it is absent in most major crop species (Underwood & Mercier, 2022).

Due to the ability of apomicts to produce seeds that retain the maternal genotype, apomixis has long been considered for applications in plant breeding (Hanna and Bashaw, 1987). Introduction of apomixis from natural species to sexual relatives by plant breeding has generally proved inefficient or unsuccessful with respect to clonal seed production (Hanna and Bashaw, 1987; Leblanc et al., 1996; Richards, 2003).

Engineered apomixis would be particularly valuable in naturally self-pollinating species where large-scale hybrid seed production is challenging. Its impact on the production of cross-pollinated crops would economize hybrid seed production, and smallholder farmers would benefit from the availability of previously inaccessible hybrid seed.

Strategies have been borrowed from mechanisms in natural apomictic species for engineering apomixis into crop plants. Two broad categories of apomictic mechanisms are sporophytic and gametophytic apomixis (Hand & Koltunow, 2014). Most efforts to date have focused on mimicking the mechanism of gametophytic apomixis where meiotic avoidance is combined with parthenogenesis to produce unrecombined, unreduced gametes and fertilization-independent embryo formation (Underwood & Mercier, 2022). Meiotic avoidance, or apomeiosis, can be achieved through mutagenesis to induce the *Mitosis instead of Meiosis*, or *MiMe* phenotype (d’Erfurth et al., 2009). *MiMe* was first implemented in Arabidopsis by combining the loss-of-function mutants *Atspo11-1*, *Atrec8* and *Atosd1* (d’Erfurth et al., 2009). In *MiMe*, the *spo11* mutation disrupts homologous recombination, and when combined with the *rec8* mutation, functionally converts meiosis I into a mitosis-like division with separation of sister chromatids and without crossing over. Then, the *osd1* mutation functions to eliminate meiosis II (d’Erfurth et al., 2009), successfully replacing meiosis with mitosis. The replacement of meiosis with mitosis using orthologous gene mutations was subsequently achieved in rice (Mieulet et al., 2016).

The capability to induce parthenogenesis is possible by utilizing members of AP2 transcription factor genes, functionally referred to as the *BABYBOOM* (*BBM*) family, first shown to induce embryos and cotyledon-like structures on seedlings in Arabidopsis (Boutillier et al., 2002). The natural apomict C*enchrus squamulatus* (syn. *Pennisetum squamulatum*), utilizes the apospory-specific genomic region *BABYBOOM-LIKE2* gene (*ASGR-BBML2*) to induce parthenogenesis. The *ASGR-BBML2* gene is expressed in the egg cell up to two days before anthesis leading to embryonic cell divisions and parthenogenetic embryo formation in the ovule (Conner et al., 2015). Furthermore, *ASGR-BBML*2 directed to the egg cell can induce haploid embryo production in sexual pearl millet (Conner et al., 2015), rice, and maize (Conner et al., 2017), pointing to the utility of this gene as a component in engineering apomixis in cereal grains. *MATRILINEAL* (*MTL*)/*NOT LIKE DAD* (*NLD*)/*ZmPLA1* also has been identified as a locus responsible for haploid seed formation in maize (Gilles et al., 2017; Kelliher et al., 2017; Liu et al., 2017), and the *Taraxacum officinale PARTHENOGENESIS* has been identified to induce haploid embryo-like structures when expressed in lettuce (Underwood et al., 2022).

By combining CRISPR/Cas9 knockout of the *MiMe* components, together with parthenogenesis induced by either *BBM, MTL or ToPAR* expression, synthetic apomixis has been successfully engineered in rice (Dan et al., 2024; Huang et al., 2025; Khanday et al., 2019; Liu et al., 2023; Song et al., 2024; Vernet et al., 2022; Wang et al., 2019; Wei et al., 2023; Xie et al., 2019; Song et al., 2024). In Arabidopsis, where parthenogenetic embryogenesis cannot be induced through *BBM* expression, clonal seeds have instead been engineered by combining *MiMe* with genome elimination after fertilization (Marimuthu et al., 2011; Ravi & Chan, 2010).

While these studies prove that apomixis is possible, additional work is needed to improve the efficiency of clonal seed production. Seed yield comparable to sexually produced hybrids and near 100% penetrance of clonal seed would be required to create a commercially viable hybrid seed product (Heidemann et al., 2025; Underwood & Mercier, 2022).

While synthetic apomixis has been successfully induced in rice and Arabidopsis, other plant species where clonal seed production could provide agronomic advantages have so far been unsuccessful at scale. Sorghum (*Sorghum bicolor*) is a cereal grain that was domesticated in sub-Saharan Africa (de Wet, 1978). It remains a subsistence food crop for millions of sub-Saharan African smallholder farmers. It is also grown worldwide for human food, livestock feed, biofuel, and forage (Hossain et al., 2022; Khaalifa & Eltahir, 2023; Kazungu et al., 2023). Compared with other major cereal crops, sorghum has increased resilience and performance under stressful conditions and on marginal land (Khalifa & Eltahir, 2023). Sorghum is naturally self-pollinating, making hybrid seed production an under-utilized source of genetic gain in resource-limited areas of the world. Engineering self-reproducing (SR) hybrids through apomixis would provide smallholder farmers with saveable, high-yielding hybrid seed that harnesses heterosis. It would also economize hybrid sorghum production and advance new hybrid varieties for specific agroecological regions in both commercial and resource-limited contexts.

Here we have demonstrated that the induction of *MiMe* via gene editing of endogenous sorghum genes *Spo11-1*, *Rec8* and two *Osd1-Like* loci in combination with parthenogenesis elicited by transgenic introduction of the *Cenchrus ASGR-BBML2* gene induces apomictic reproduction in sorghum. Induction of SR hybrids in two genetic backgrounds using two different methods were followed through multiple generations and clearly demonstrate the inheritance of the hybrid genotype. We observed a significant reduction in seed number per plant compared to the wild-type conventional hybrid, with clear evidence that *Osd1-Like* gene dosage is one factor crucial to plant growth and seed set in induced apomictic sorghum hybrids.

## Results

### Validating *Mitosis Instead of Meiosis* (*MiMe*) component function in the Tx430 variety of sorghum

The progression of meiosis in developing meiocytes in wild-type sorghum are shown in Figure 1A-D and mature tetrads are shown in Supplemental Figure 1. *Spo11-1* was targeted by CRISPR/Cas9 gene editing to prevent double strand breaks and chromosome pairing at meiosis I (Supplemental Table 1). Cytology of male meiocytes in *spo11-1* mutant plants (Figure 1 E-H) showed modifications to meiosis with 20 univalents (Figure 1 F, G) instead of 10 bivalents observed in wild-type TX430 (Figure 1B, C). Random segregation of chromosomes occurred during meiosis I (Figure 1H), producing unbalanced tetrads, polyads and triads (Supplemental Figure 1) at the completion of meiosis II. The *Spo11-1* mutant plants were almost completely sterile following self-pollination.

**Figure 1:**
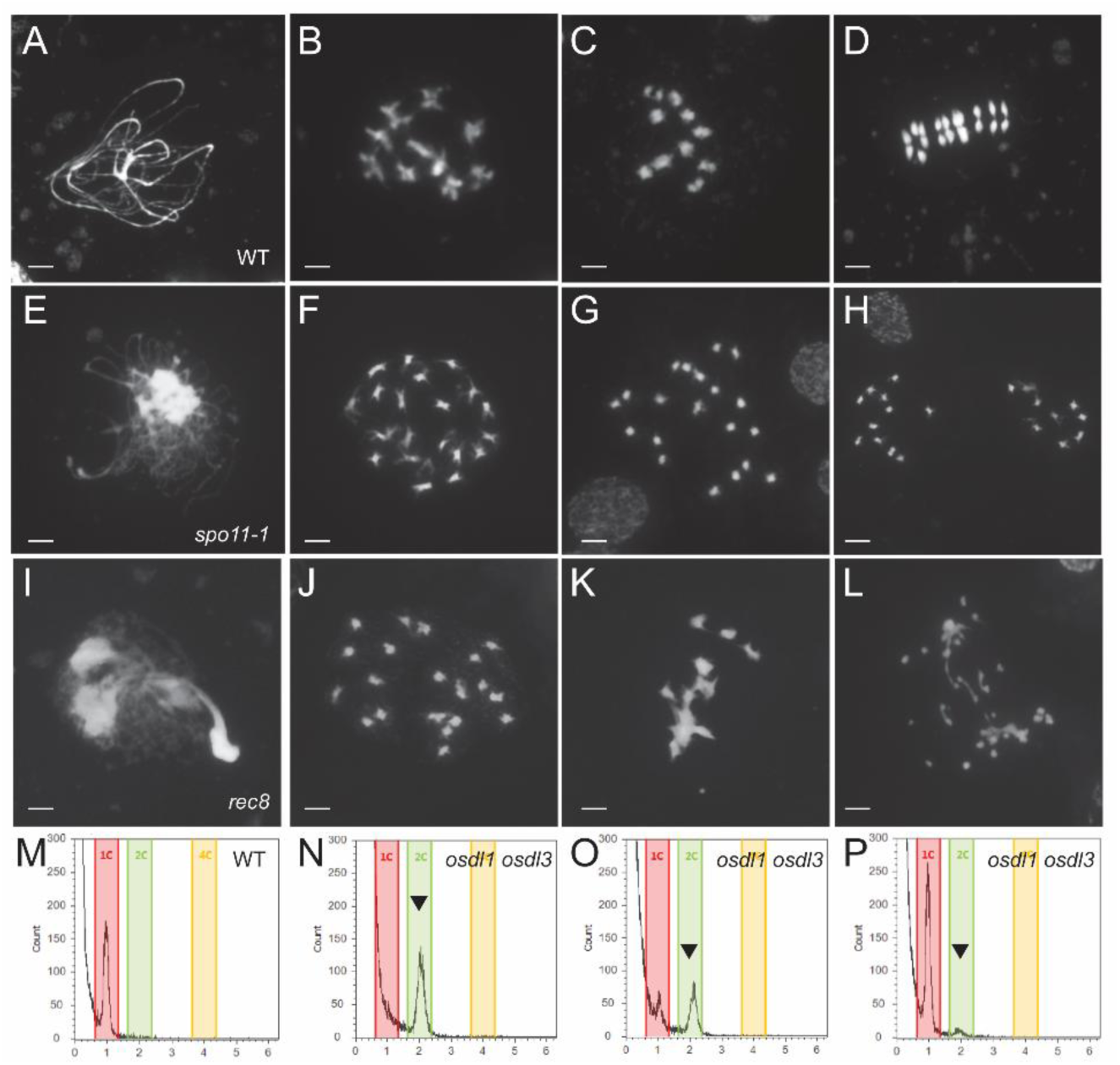
Male meiosis in sorghum. (A-D) WT Tx430 control, (E-H) *spo11-1* mutant, (I-L) *rec8* mutant; at pachytene stage (A, E, I), diakinesis (B, F, J), early metaphase I (C, G, K), early anaphase I (D, H, L). Bars = 50 µm. (M-P) Flow cytometry histograms of stained nuclei isolated from pollen of WT control (M), and plants containing four out-of-frame *OsdL1*/*OsdL3* alleles (N), three out-of-frame and one in-frame *OsdL1*/*OsdL3* alleles (O) and one out-of-frame and three in-frame OsdL1/OsdL3 alleles (P). Arrowhead indicates diploid pollen nuclei (relative propidium iodide fluorescence X axis, number of nuclei Y axis).

*Rec8* was targeted for CRISPR/Cas9 knockout to prevent sister chromatid cohesion at meiosis I (Supplemental Table 1). Cytology of male meiocytes in *rec8* mutant plants (Figure 1I-L) showed reduced homolog pairing and 20 univalents (Figure 1I, J) instead of 10 bivalents (Figure 1B, C). Early sister chromatid segregation and chromosome fragmentation occurred during meiosis I (Figure 1K, L) and polyads at completion of meiosis II (Supplemental Figure 1). Self-pollinated *Rec8* mutant plants were completely sterile and did not produce any seed.

Sorghum contains two tightly linked *Osd1-Like* genes (*OsdL1* and *OsdL3*) separated by 7802 bp rather than the single *Osd1* gene identified and functionally characterized in rice (Mieulet et al., 2016). A third *Osd1*-*Like* gene was also identified in sorghum, but it was outside the *Osd1* clade identified in rice (Mieulet et al., 2016). The function of each of these putative *OsdL1* and *OsdL3* homologs in the progression of meiosis was examined by individually targeting each gene for knockout with CRISPR/Cas9 (Supplemental Table 1). *Osd* mutations are expected to produce unreduced diploid gametes, and when self-fertilized, these mutants should generate tetraploid progeny. Knockouts in either *OsdL1* or *OsdL3* however, only produced diploid progeny (Table 1). Heterozygous alleles of *OsdL1* and *OsdL3* single gene edits linked in *trans* also produced diploid progeny (Table 1). Collectively, these results indicate that a mutation in each individual *OsdL* gene, either *OsdL1* or *OsdL3,* is insufficient to give a meiotic non-reduction phenotype.

**Table 1.**
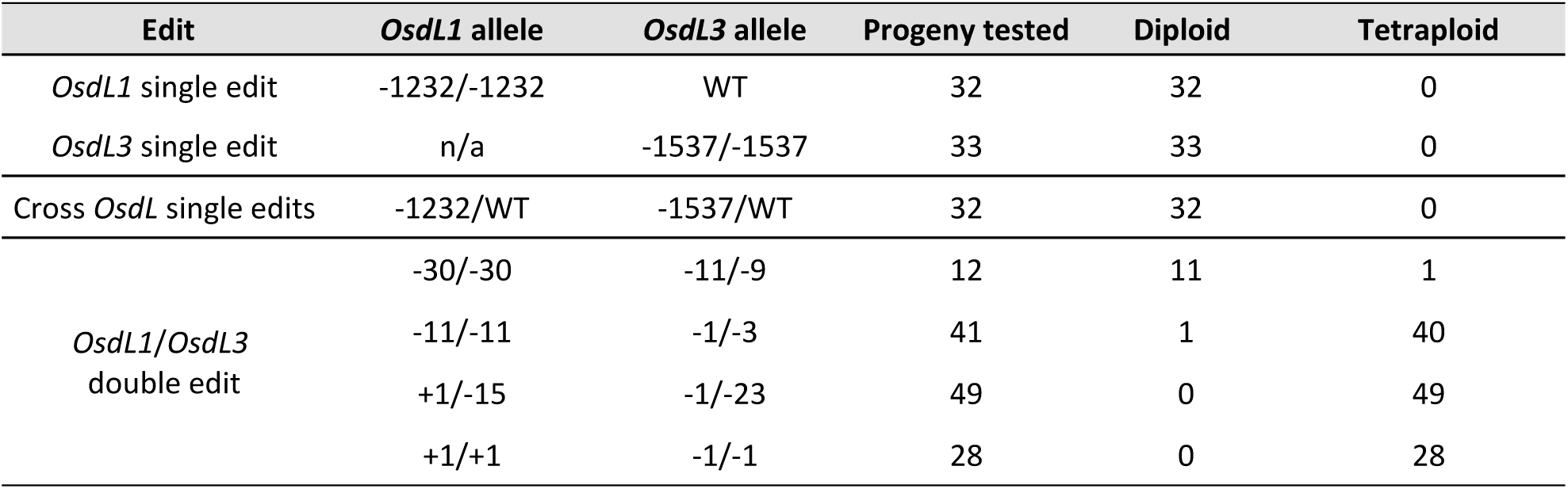
Ploidy of progeny from *OsdL* edited plants.

To examine the possibility of functional redundancy, both *OsdL1* and *OsdL3* genes were targeted with CRISPR/Cas9 to knockout gene function (Supplemental Table 1). Double knockout mutants were obtained with four frameshift mutations. In addition, plants with three frameshift mutations and one in-frame mutation at either the *OsdL1* or the *OsdL3* loci were also obtained. Pollen flow cytometry analyses (Figure 1 M-P) showed that haploid pollen was produced in wild-type Tx430 (Figure 1M), whereas double knockout plants produced diploid pollen (Figure 1N). Plants containing three out-of-frame alleles and one in-frame allele produced both haploid and diploid pollen (Figure 1O), whereas plants containing one in-frame allele and three out-of-frame alleles produced mostly haploid pollen (Figure 1P).

To characterize the combined effect of both male and female gamete non-reduction, we measured ploidy levels of the progeny. All progeny from the double knockout plants were tetraploid (Table 1). Progeny from a parent that contained three out-of-frame and one in-frame alleles were mostly tetraploid, whereas progeny from a parent containing one in-frame allele and three out-of-frame alleles were mostly diploid (Table 1). Collectively these data indicate that the *OsdL1* and *OsdL3* genes are functionally redundant, and that mutations in both genes are required to give a non-reduction phenotype.

### Parthenogenesis induction in the sorghum variety Tx430

To examine parthenogenesis induction in sorghum, six constructs were initially prepared (Supplemental Table 2). Two constructs contained the previously identified apomictic *Cenchrus ASGR-BBML2* promoter combined with either the *ASGR-BBML2* genomic or the coding gene sequence (Conner et al., 2017). The expression pattern of the *ASGR-BBML2* promoter was unknown in sorghum, therefore, an additional four constructs were developed that contained the maize orthologous promoters of the originally identified Arabidopsis *DD45* and *RKD2* egg cell preferred expressed genes (Albertsen et al., 2013), fused to the *Cenchrus ASGR-BBML2* genomic or coding gene sequence (Conner et al., 2017).

Parthenogenetic events were evaluated by cytological observations of embryo development in the female embryo sac post-anthesis in wild-type and transgenic plants. Prevention of self-pollination in a wild-type Tx430 sorghum plant results in the female gametophyte remaining unfertilized with a single egg cell and two unfused polar nuclei (Figure 2A-C). After double fertilization in Tx430, the embryo develops from the egg fertilized by one sperm cell, and the endosperm develops following fusion of the second sperm cell with the two polar nuclei (Figure 2D). In a hemizygous transgenic plant containing an egg cell targeted *ASGR-BBML2* transgene, a 1:1 segregation and inheritance of the transgene is expected with parthenogenesis occurring in 50% of the female gametes.

**Figure 2.**
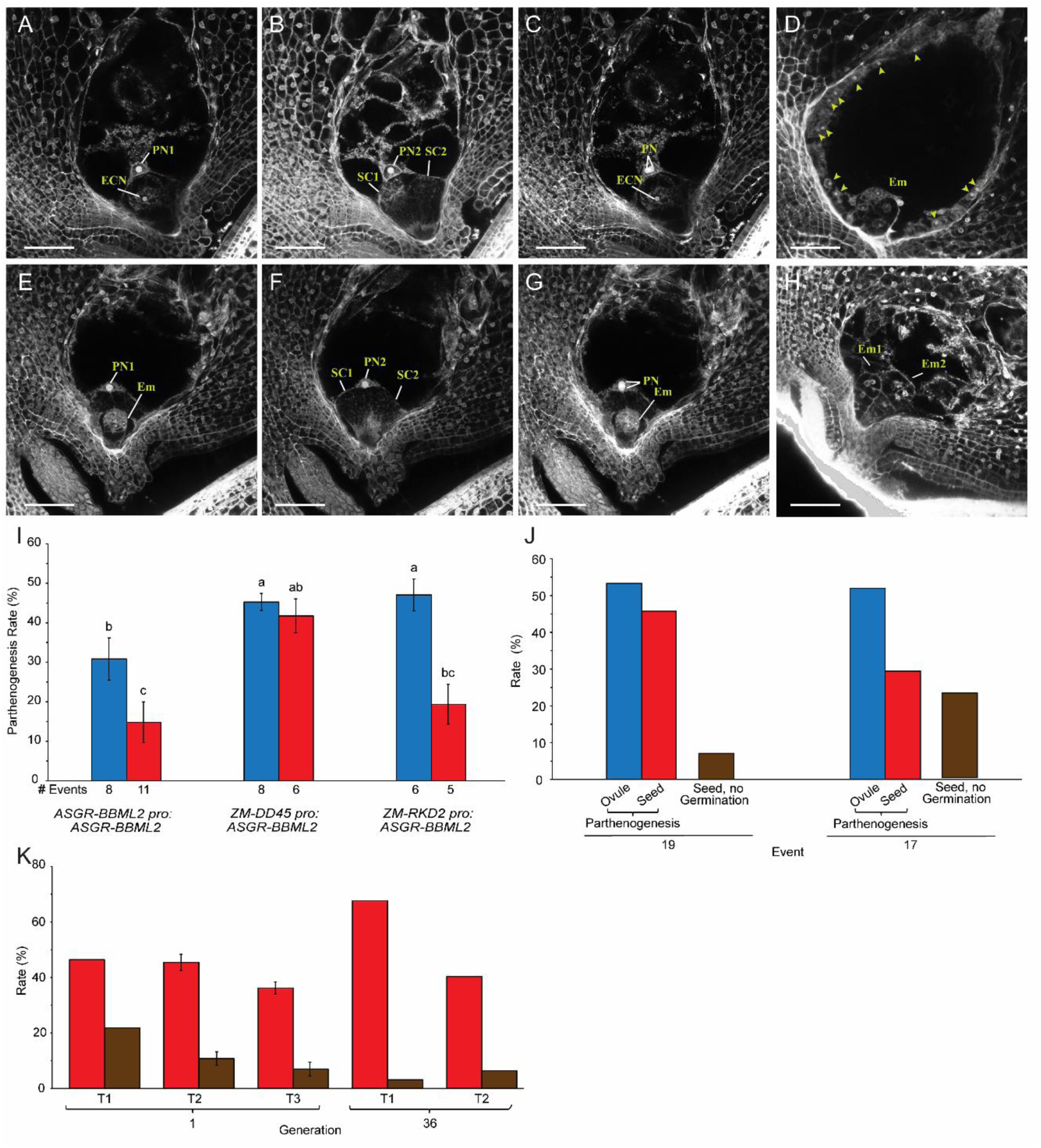
Characterization of parthenogenesis in sorghum. (A-C) Two focal planes (A, B) of an emasculated embryo sac of WT Tx430 control, with an unfertilized egg cell nucleus (A), two unfused polar nuclei (A,B) and two synergid cells (B), with merged image in (C). (D) Fertilized embryo sac ∼2 days after pollination of a WT Tx430 control, with a fertilized embryo and syncytial endosperm nuclei (arrowheads) (E-G) Two focal planes (E, F) of an emasculated embryo sac with a parthenogenetic embryo (E), two unfused polar nuclei (E, F) and two synergid cells (F), with merged image in (G). Bars = 45 um. (H) An emasculated embryo sac with two parthenogenetic embryos. (I) Mean parthenogenesis rate in constructs of *ASGR-BBML2* genomic (blue) and cds sequences (red) driven by three different promoters (*ASGR-BBML2*, maize *DD45* and maize *RKD2* promoters), with number of events per construct shown. Constructs without significant differences (P>0.05) are labeled with the same letters. Similar rates (P>0.05 from two-tailed pairwise comparisons) are labeled with the same letter. (J) Comparison of parthenogenesis rate determined by cytological analysis (blue), progeny flow cytometry (red), and seed non-germination rates (brown) observed in events 19 and 17 of *ASGR-BBML2 pro:ASGR-BBML2* (PHP83000). (K) Comparison of parthenogenesis rate determined by flow cytometry (red) and seed non-germination rate (brown) of events 1 and 36 of *ZM-DD45 pro:ASGR-BBML2* cDNA (PHP94292) in T1, T2 and T3 generations.

From the transformation of the six constructs a total of 44 unfertilized hemizygous transgenic T0 plants, with a range of 5-11 T0 plants for each construct, were cytologically examined for parthenogenesis (Figure 2I). This was determined by the development of an embryo-like structure at the micropylar end of the embryo sac, the presence of two unfused polar nuclei, and the corresponding absence of endosperm development (Figure 2E-G). The development of two parthenogenetic embryos was observed in a single female gametophyte in a small number of ovules (Figure 2H). Three of the six constructs, containing the maize *DD45* promoter fused to the *ASGR-BBML2* gDNA or cDNA, or the *RKD2* promoter fused to the *ASGR-BBML2* gDNA, exhibited average parthenogenesis rates greater than 40% (Figure 2I). The other three constructs, containing the native *ASGR-BBML2* promoter fused to the *ASGR-BBML2* gDNA or cDNA, or the *RKD2* promoter fused to the *ASGR-BBML2* cDNA, exhibited an average parthenogenesis rate that ranged from 10-30% (Figure 2I). This suggested that the maize *DD45* promoter induced parthenogenesis at equivalent rates for both the *ASGR*-*BBML2* genomic and cDNA sequences. But the native *ASGR-BBML2* and the maize *RKD2* promoters were less efficient in stimulating parthenogenesis when linked to the cDNA relative to the genomic coding sequence. Position effects of transgene insertion in the genome on parthenogenesis frequency, cannot be ruled out.

Two independent T1 progeny events containing the native *ASGR-BBML2* promoter fused to the gDNA were examined by cytology and flow cytometry to examine rates of embryo initiation and germination of seeds containing haploid embryos. In event 19 the cytology and flow cytometry showed similar rates of haploid embryo initiation and development with 6.3% non-germinated seed, whereas in event 17, the frequency of haploid embryo initiation was much greater than observed haploid progeny with 20.3 % non-germinated seed (Figure 2J). These results suggest that position effects may also influence the maturation of haploid embryos in seeds.

For forward combination of parthenogenesis induction with *MiMe*, another parthenogenesis construct was designed to generate transgenic lines containing the maize *DD45* promoter and *Cenchrus ASGR-BBML2* coding sequence, which had demonstrated good penetrance of the parthenogenesis phenotype (Figure2I). This construct was designed to produce plants that would not contain additional T-DNA components (Supplemental Table 2). An advantage of this construct design is that it should eliminate the effect of adjacent promoters in the T-DNA from potentially influencing the expression pattern or level of the parthenogenesis cassette. As shown in Figure 2K, flow cytometry of two transgenic events containing this construct, showed a high frequency of haploid progeny across multiple generations. Two embryos in a single seed were observed in 3.0% of progeny seed across both events.

Germination and subsequent ploidy determination of these dual embryo progeny seed revealed that most dual embryos were haploid-haploid pairs and rarely haploid-diploid pairs (data not shown). It is unknown if the dual haploid embryos originate by embryo budding or possibly synergid embryo induction and if the diploids in the haploid pairs arise from sexual fertilization or by spontaneous doubling as occurs in some species (Fomicheva et al., 2024).

### Two-step induction of synthetic apomixis in a Tx623/Tx430 hybrid

Two different approaches were used to combine *MiMe* with parthenogenesis to produce sorghum SR hybrids. A two-step approach was used in the generation of SR Tx623/Tx430 sorghum hybrids. In the first step, a characterized Tx430 transgenic line with the maize *DD45* promoter*:ASGR-BBML2* coding gene sequences was crossed with a wild-type Tx623 sorghum line to generate a F1 hybrid. In the second step, immature embryos from this cross were used in a secondary transformation to introduce CRISPR/Cas9 together with *MiMe* gRNAs (Figure 3A). The frequency of editing at each individual *MiMe* locus was consistent and ranged from 45.3.% to 47.3% (Supplemental Table 3). In total, 33 plants were retained with a hemizygous *ASGR-BBML2* gene and a combination of unique edits at the *MiMe* loci. T0 biallelic edits were germinal and were clonally inherited in T1 progeny (Supplemental Table 3).

**Figure 3.**
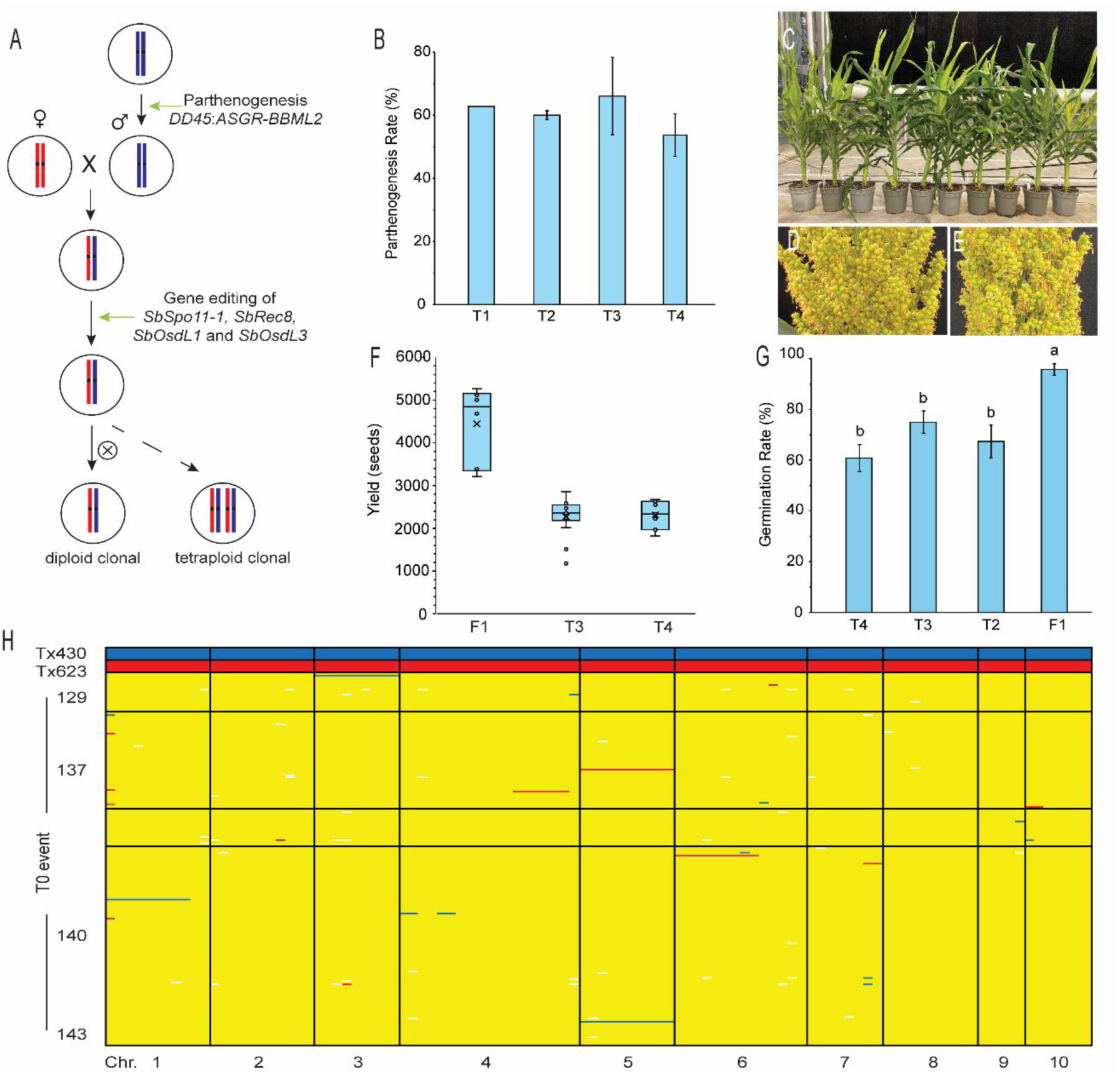
Two-step synthetic apomixis in Tx623/Tx430 hybrid sorghum. (A) Schematic representation of two-step induction of synthetic apomixis. (B) Average parthenogenesis rate of T1, T2, T3 and T4 generations of event 143, +/-se. (C-E) Comparison of plant and panicle morphology. (C) Whole plant phenotype of synthetic apomicts and F1 sexual control (indicated with *). Panicle of F1 sexual hybrid (D), and apomictic T2 plant (E). (F) Mean seed yield per plant and yield variability in F1 hybrid control and T3 and T4 generations. (G) Mean seed germination rates F1 hybrid control and T2, T3 and T4 generations of event 143, +/-se. (H) SNP marker analysis of the Tx623 and Tx430 parental controls and 238 T1 progeny from four selected T0 events. The Tx430 alleles are marked in blue, Tx623 alleles in red, heterozygous alleles in yellow, and missing marker calls in white.

A range of phenotypes were observed in T0 plants in the greenhouse, some plants showed significantly delayed and aberrant development with a reduction in plant height, leaf and stem curling, and reduced seed set (Supplemental Figure 2). It is noteworthy that seed number per plant was significantly reduced in the T0 apomictic hybrids with complete knockouts of all four *OsdL1* and *OsdL3* alleles (Table 2). There was, however, a clear improvement in the fertility of T0 plants that contained three out-of-frame alleles and one in-frame allele in the *OsdL1* and *OsdL3* loci relative to T0 plants that contained four out-of-frame alleles (Table 2). Four independent T0 events were chosen for further characterization of apomictic progeny in the T1 generation relative to a hybrid control. Multiple progeny from each event, however, showed reduced seed set in comparison to the sexual hybrid control (Table 3).

**Table 2.**
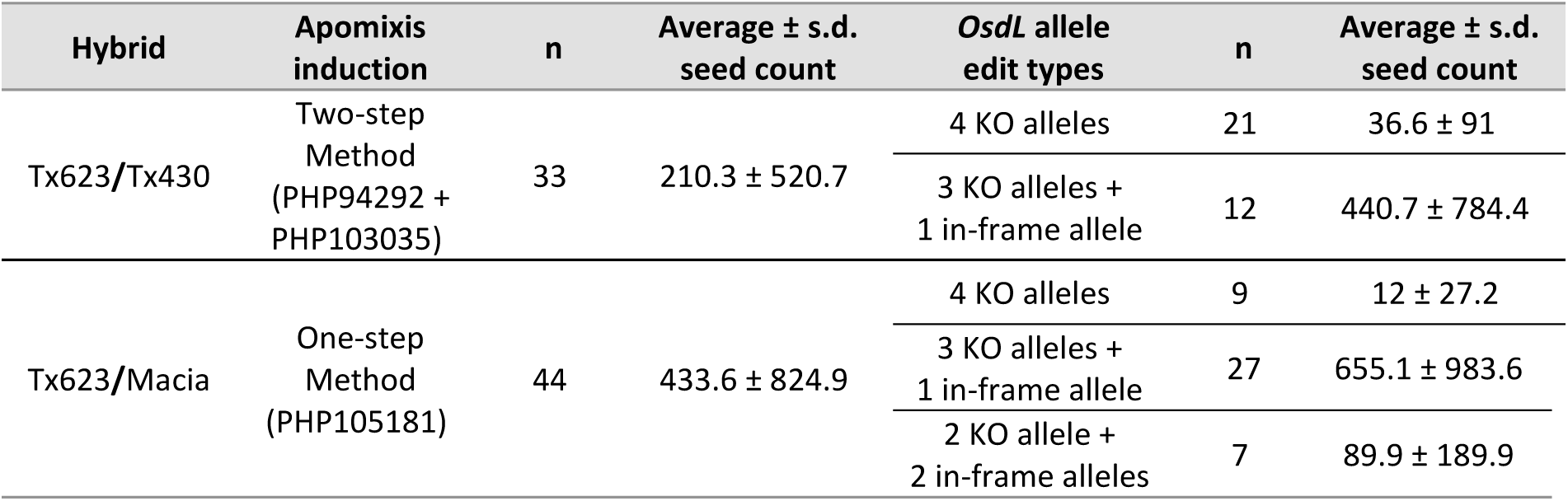
Effect of *OsdL* gene dosage on T1 seed production in Tx623/Tx430 and Tx623/Macia apomictic plants. KO is knockout.

| Hybrid | Apomixis induction | n | Average ± s.d. seed count | *OsdL* allele edit types | n | Average ± s.d. seed count |
| --- | --- | --- | --- | --- | --- | --- |
| Tx623/Tx430 | Two-step Method (PHP94292 + PHP103035) | 33 | 210.3 ± 520.7 | 4 KO alleles | 21 | 36.6 ± 91 |
| | | | | 3 KO alleles + 1 in-frame allele | 12 | 440.7 ± 784.4 |
| Tx623/Macia | One-step Method (PHP105181) | 44 | 433.6 ± 824.9 | 4 KO alleles | 9 | 12 ± 27.2 |
| | | | | 3 KO alleles + 1 in-frame allele | 27 | 655.1 ± 983.6 |
| | | | | 2 KO allele + 2 in-frame alleles | 7 | 89.9 ± 189.9 |

**Table 3.** Analysis of progeny ploidy and seed set in selected Tx623/Tx430 and Tx623/Macia apomictic events. n.t. not tested.

| Hybrid | Apomixis induction | T0 event | T1 seed count | Diploid frequency (%) | T1 plants (n) | T2 seed count (average ± s.d.) |
| --- | --- | --- | --- | --- | --- | --- |
| Tx623/Tx430 | Two-step Method<br>(PHP94292 + PHP103035) | 140 | 202 | 87.5% | 6 | 68.5 ± 61.6 |
| | | 143 | 201 | 74.8% | 4 | 1264.8 ± 222.5 |
| | | 137 | 449 | 82.3% | 13 | 300.8 ± 258.4 |
| | | 129 | 491 | 68.0% | 3 | 601.7 ± 382.5 |
| | F1 control | n/a | n/a | n/a | 6 | 4222 ± 329.7 |
| Tx623/Macia | One-step Method<br>(PHP105181) | 105 | 559 | 48.9% | n.t. | n.t. |
| | | 120 | 3533 | 29.9% | 6 | 1488.7 ± 537.8 |
| | | 60 | 1383 | 53.8% | 2 | 195 ± 11.3 |
|  |  | 78 | 2042 | 57.5% | n.t. | n.t. |
| | F1 control | n/a | n/a | n/a | 15 | 3860 ± 1678 |

The Tx430 *ASGR-BBML2* parthenogenesis event produced 43% haploid progeny when hemizygous in the sexual parent plant (Figure 2K). When this parthenogenesis event was combined with *MiMe* mutations with three knockouts in both *OsdL* genes and one in-frame *OsdL* allele to produce a synthetic apomictic hybrid parent, the expected progeny ratios were 86% parthenogenetic diploids and 14% non-parthenogenetic tetraploids. The observed frequency of diploid progeny ranged from 68.0-87.5% across the 4 independent events (Table 3).

To confirm the successful production of synthetic apomictic clonal sorghum progeny, a set of 104 polymorphic SNP markers across the 10 chromosomes of sorghum was used for analyses. In total, 238 T1 progeny were analyzed that were produced from four independent T0 events, including 184 diploid individuals and 54 tetraploid individuals. Over 96% of the analyzed T1 progeny (229 of the 238 individuals) were heterozygous on all chromosomes and clonal (Figure 3H). Therefore, the identification of hybrid diploid clonal progeny demonstrates that the combination of *MiMe* and *ASGR-BBML2* was successful in the female gamete to induce synthetic apomixis. The identification of hybrid tetraploid progeny demonstrates that *MiMe* was active in both the male and female gametes. Unexpectedly, nine individual progeny showed a SNP allele pattern that suggested these were aneuploid. Among these putative aneuploid progeny, three individuals had lost one copy of a single chromosome, and six individuals had a partial deletion of a single chromosome (Figure 3H).

One event (event 143) that contained one in-frame and one out-of-frame allele in *OsdL1* and two out-of-frame alleles in *OsdL3* and showed good seed set in the T1 generation was selected for further analysis in subsequent T2, T3 and T4 generations. The relative parthenogenesis frequency as measured by the frequency of diploid individuals in the T2, T3 and T4 generations was stable across generations (Figure 3B). The maintenance of heterozygosity in both the diploid and tetraploid progeny was evaluated in each generation using the same set of 104 polymorphic SNP markers described above. From the 207 T2 generation individual progeny, 638 T3 generation progeny and 126 T4 generation progeny, we observed that the *MiMe* phenotype is stable and highly penetrant across generations. Similar to the results observed in the T1 generation, over 98% of the T2 progeny, 93% of the T3 progeny and 97% of the T4 progeny were clonal, with SNP markers showing heterozygous alleles across all chromosomes (Supplemental Figure 3). The remaining individuals in each generation were aneuploid with either a large deletion of a single chromosomal segment or a deletion of an entire single chromosome.

In addition to molecular analysis of event 143, whole plant phenotype observations showed no differences between apomictic individuals and the hybrid controls during vegetative development (Figure 3C). Differences were noted, however, during panicle development where many florets did not show evidence of seed development, which correlated with reduced seed set (Figure 3D-F). The relative reduction of seed set was consistent across multiple generations (Figure 3F). Seeds were phenotypically normal (Supplemental Figure 4), although seed germination of the apomictic hybrids also was reduced relative to the hybrid controls (Figure 3G). Flow cytometry showed that the ploidy of the endosperm of the synthetic apomictic lines was increased to 6n, compared to 3n endosperm in the sexual control (Supplemental Figure 5). This is expected given the presence of two diploid central cell nuclei fertilized by a diploid sperm cell nucleus. Whether this increase in endosperm ploidy has an influence on germination and positive or negative consequences on seed quality is yet to be determined. These data indicated that SR hybrid sorghum can be generated in a two-step induction approach and the induced synthetic apomictic seed set is maintained over four generations.

### One-Step transformation method for induction of synthetic apomixis in sorghum in a Tx623/Macia hybrid

A second approach to induce synthetic apomixis in sorghum was conducted in a hybrid cross between Tx623 as the female parent and the African variety Macia. Immature embryos from the F1 hybrid cross were used in a primary transformation to introduce the maize *DD45* promoter*:ASGR-BBML2* coding sequence parthenogenesis cassette together with CRISPR/Cas9 and *MiMe* gRNAs in a single T-DNA construct (Figure 4A). The efficiency of CRISPR/Cas9 mediated mutations at each of the four target loci was evaluated by amplicon sequencing and was found to be consistent, ranging from 51.3% to 53.1% (Supplemental Table 3). In total, 44 plants were retained and analyzed for *MiMe* loci edits (Supplemental Table 4) and for inheritance of *ASGR-BBML2*. Similar to the two-step synthetic apomixis experiment, it was clear that there was a relative improvement in the overall reproductive fertility in the T0 plants that contained 3 out-of-frame alleles and 1 in-frame allele in the *OsdL1* and *OsdL3* loci relative to the T0 plants that contained 4 out-of-frame alleles (Table 2).

**Figure 4.**
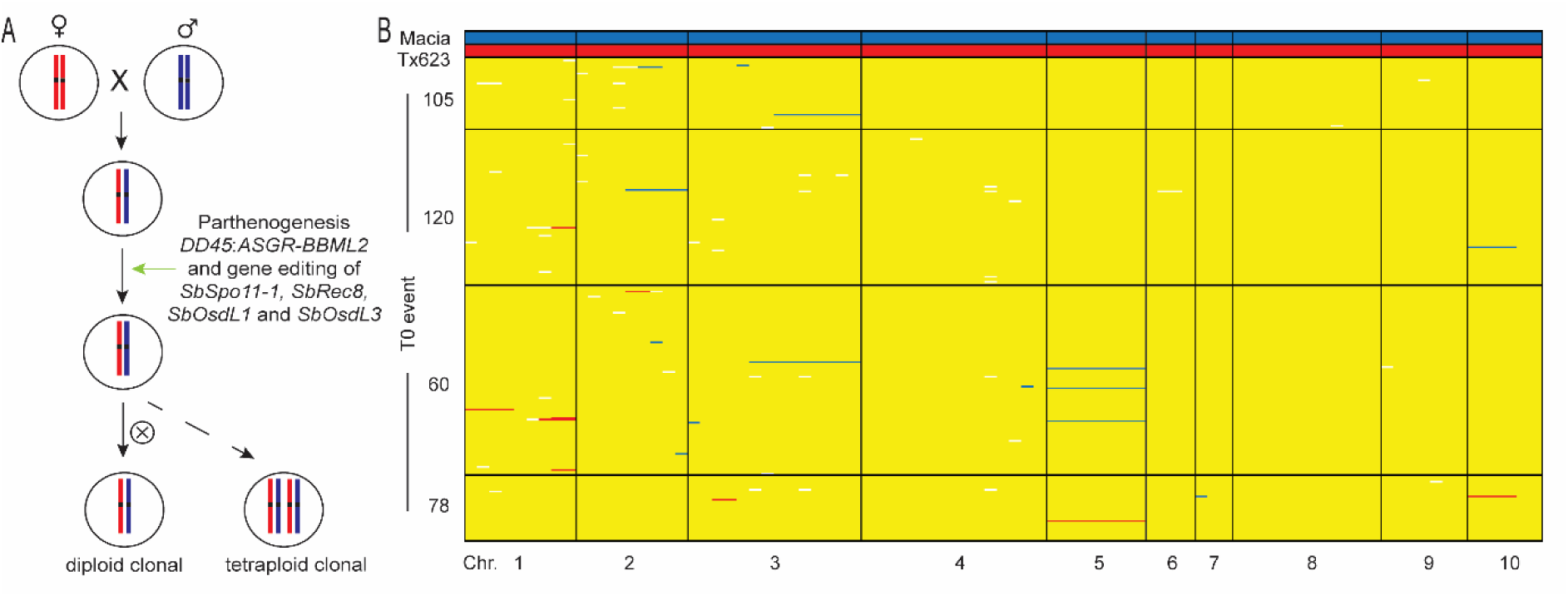
One-step synthetic apomixis in Tx623/Macia hybrid sorghum. (A) Schematic representation of one-step induction of synthetic apomixis. (B) SNP marker analysis of the Tx623 and Macia parental controls and 295 T1 progeny from four selected T0 events. The Macia alleles are marked in blue, Tx623 alleles in red, heterozygous alleles in yellow, and missing marker calls in white.

Four independent T0 events were chosen for further characterization of apomictic T1 progeny relative to a hybrid control. Seed set of the T1 individuals was reduced and variable relative to the hybrid control (Table 3). Parthenogenesis frequencies in T1 progeny of these selected apomictic events ranged from 29.9% to 57.5% (Table 3). A set of 87 polymorphic SNP markers was used to evaluate maintenance of heterozygosity. In total, 295 T1 progeny were analyzed, including 137 diploid individuals and 158 tetraploid individuals. Similar to the *MiMe* results shown above for the Tx623/Tx430 hybrid, over 96% of the T1 progeny analyzed were clonal (284 of the 295 individuals), with heterozygous alleles across all chromosomes (Figure 4B).

Only 11 individuals showed large or entire chromosomal deletions that suggested the production of aneuploid progeny (Figure 4B). These data indicated that SR hybrid sorghum can also be generated when all of the induction components are introduced within one T-DNA plasmid.

## Discussion

This is the first report of synthetic apomixis induction in hybrid sorghum and demonstration that the resultant self-reproducing hybrids to maintain heterozygosity in multiple self-fertilized progeny generations. Introduction of synthetic apomixis into hybrid sorghum was successful by combining CRISPR/Cas9 induced mutations in the *MiMe* loci together with an egg cell expressed *ASGR-BBML2* transgenic component.

Importantly, synthetic apomixis was demonstrated in two hybrid backgrounds using two different induction approaches (one-step and two-step integration techniques). The two-step approach has the advantage of first producing stable parthenogenesis phenotypes in all progeny, whereas the one-step approach minimizes the number of transformation steps but produces more variability in parthenogenesis.

Contrary to previous reports in rice (Khanday et al., 2019, Vernet et al., 2022) that showed an improvement in the induction of parthenogenesis and resulting synthetic apomictic progeny when using a one-step technique, in sorghum we observed greater variability in parthenogenesis with a similar approach. Nevertheless, both approaches offer unique advantages to implement apomixis. Identifying a construct design that consistently delivers high rates of parthenogenesis is essential.

We demonstrate parthenogenesis induction rates of approximately 40% in multiple independent transgenic lines and show that the use of a maize egg cell *DD45* promoter with the *Cenchrus ASGR-BBML2* coding sequence induced a more penetrant parthenogenesis phenotype than the *ASGR-BBML2* promoter in sorghum. In addition, the observed event to event variation of parthenogenesis suggested that somaclonal variation associated with the transformation process or positional effect of the transgene integration site may be influencing the penetrance and expressivity of the parthenogenesis phenotype. Furthermore, our observation of the development and ploidy of dual embryos may highlight the importance of the developmental timing and specificity of the egg cell promoter driving the *ASGR-BBML2* parthenogenesis gene.

Progeny seed germination was reduced in some transgenic lines containing solely the maize *DD45* promoter:*ASGR*-*BBML2* construct. This suggests that the initiation of embryogenesis through parthenogenesis may have additional deleterious effects on the development of the embryo and seed. Improvement of parthenogenesis induction may benefit from further refinement of egg cell specific expression of *ASGR*-*BBML2*, as well as the addition of other parthenogenesis genes that may supplement *ASGR-BBML2* activity (Ren et al., 2024, Underwood et al., 2022, Huang et al., 2025).

In both synthetic apomixis experiments, we observed 96.2% of all progeny maintained complete heterozygosity. In the two-step apomictic approach, incomplete penetrance of parthenogenesis results in 77.3% diploid and 22.7% tetraploid progeny. Thus, 73.9% of the total progeny are true clonal progeny that are diploid and heterozygous. In the one-step apomictic approach, penetrance of parthenogenesis was more variable with 46.4% diploid and 53.6% tetraploid progeny, resulting in 43.4% of the total progeny as true clonal progeny. Finally, the remaining 3.8% of all progeny showed evidence of aneuploidy. Marker analysis of these individuals showed a large, contiguous deletion of either a portion of a chromosome or an entire chromosome, suggesting that at a low frequency chromosome fragmentation or mis-segregation may be occurring in this *MiMe* system (*spo11-1*, *rec8*, *osdL1*, *osdL3*). Similarly, previous studies have also identified the production of aneuploid progeny in *MiMe*-2 (*spo11-1, rec8, tam*) genotypes in *Arabidopsis* and tomato (d’Erfurth et al., 2009; Wang et al., 2024; Liu et al., 2023).

In Arabidopsis and rice, a single *Osd1* gene is sufficient to promote meiotic cell cycle progression, and mutations in this gene result in unreduced male and female gametes (d’Erfurth et al., 2009; Mieulet et al., 2016). However, incomplete penetrance of the non-reduction phenotype resulted in reduced seed set due to male/female gamete ploidy imbalance at fertilization (d’Erfurth et al., 2009; Mieulet et al., 2016). In sorghum we show that two tightly linked *Osd1-Like* genes function in meiotic progression. Although our analyses of *OsdL1* and *OsdL3* four allele knockout mutants showed high penetrance of the non-reduction phenotype in the male gametes, this allelic combination unexpectedly produced significantly less seed in the synthetic apomicts compared to the sexual hybrid control. This result suggests that incomplete non-reduction in the female gametes may be a limiting factor that influences balanced fertilization and development of progeny seed. Detailed cytological characterization of female meiosis may help understand if *OsdL1* and *OsdL3* functions equivalently in male and female meiosis.

In addition, these two genes in sorghum may be essential for additional non-meiotic cell cycle functions. In Arabidopsis, both *OSD1*/*GIG* and the related gene *UVI4* have been shown to affect mitotic cell cycle regulation (Iwata et al., 2011, Iwata et al., 2012). Our analyses of *OsdL1* and *OsdL3* mutant combinations in sorghum that contained three out-of-frame and one in-frame allele showed an incomplete non-reduction phenotype. However, this allelic combination unexpectedly led to improved seed set in apomicts compared to the four out-of-frame allelic combination, suggesting that the functional dosage of *OsdL* genes is important for production of viable gametes and subsequent viable progeny. The unique complement of edited alleles produced across events did not enable a complete evaluation of the full or partial functional redundancy of the *OsdL1* and *OsdL3* genes. Interestingly, the non-reduction component of *MiMe* was also identified as a key step for further investigation in characterization of *MiMe* genotypes in tomato (Wang et al., 2024). Therefore, *OsdL* gene dosage appears to be a critical component in enabling full fertility in this apomictic system.

The level of synthetic apomictic seed set observed in sorghum was, however, not equivalent to that of hybrid seed set in sexual controls indicating other factors are required to achieve efficient seed set. Similarly, several studies in rice have reported reduced fertility of synthetic apomictic lines (Dan et al., 2024; Huang et al., 2025; Khanday et al., 2019; Song et al., 2024; Vernet et al., 2022; Wang et al., 2019). Each of the component factors inducing synthetic apomixis in both rice and sorghum, modifies aspects of embryo and endosperm development producing a diploid embryo and a 6x endosperm instead of the typical 3x persistent seed endosperm. This could influence seed development and germination. In this study we identified that both the choice of promoter to drive ASGR-*BBML2* expression for optimal parthenogenesis and embryo development and that *OsdL* dosage for meiotic non-reduction are critical factors for optimal induction of synthetic apomixis.

In conclusion, we have shown that sorghum is a crop in which synthetic apomixis is a reality. Clonal, diploid progeny from hybrid apomictic sorghum develops similarly to hybrid controls during vegetative growth. Seed set is reduced, but stable, across generations relative to a sexual hybrid control. While additional improvements in fertility will be required for commercial grain production, this technology has the potential to lead to significant improvements in agriculture by capturing heterosis and maintaining hybrid vigor through seed in sorghum. Resolving inefficiencies in this process would be an important step for enabling field production of apomictic hybrids in sorghum.

## Materials and Methods

### Plant materials and handling

Sorghum genotypes Tx430, ATx623 and the African landrace Macia (PI 565121) were requested from USDA GRIN and were cultivated in the greenhouse with 78°F during the day and 68°F night temperatures under a 16 h photoperiod. To produce F1 embryo donor material for sorghum transformation for the two-step approach, individual crosses were performed using CMS sterile Tx623 (ATx623) as a female parent with the transgenic Tx430 (PHP94292 event 1) parthenogenesis line as the male parent. To produce F1 embryo donor material for sorghum transformation for the one-step approach, individual crosses were performed using CMS sterile Tx623 (ATx623) as a female parent with the non-transgenic Macia line as the male parent. Furthermore, fertile F1 progeny control seed of these two hybrids was produced by crossing the CMS sterile ATx623 as a female parent with either the non-transgenic Tx430 or non-transgenic Macia lines as the male parent. Crosses of Tx430 CRISPR/Cas9 edited lines were completed using hot water emasculation of the female parent immature panicle.

### Construct design

Seven plasmids were used for sorghum transformation to test parthenogenesis (Supplemental Table 2). The *Cenchrus ASGR-BBML2* promoter, *ASGR-BBML2* genomic and *ASGR-BBML2* coding sequence was generously provided by Peggy Ozias-Akins. PHP83000 and PHP82772, contained the *Cenchrus ASGR-BBML2* promoter driving the genomic or coding sequence of the *ASGR-BBML2* gene, respectively, and a PMI selectable marker. PHP86483 and PHP86482 contained the maize ZMDD45 promoter, and PHP86484 and PHP86796 contained the maize RKD2 promoter (Albertsen et al., 2013), each driving the genomic or coding sequence of the *ASGR-BBML2* gene, respectively, and a HRA selectable marker. A final plasmid, PHP94292, contained the same maize *DD45:ASGR-BBML2* cassette as PHP86482.

Six plasmids were used for sorghum transformation to test gene editing of *MiMe* gene loci (Supplemental Table 2). Plasmids PHP85484, PHP83595, PHP83596, PHP83436 each contained two gRNA cassettes targeting each gene *Sb-Spo11-1*, *Sb-Rec8*, *Sb-OsdL1*, *Sb-OsdL3*, respectively, a maize optimized Cas9 cassette, and a PMI selectable marker. Plasmids PHP85930 and PHP110251 contained two gRNA cassettes targeting the *Sb-OsdL1* and *Sb-OsdL3* genes, a maize optimized Cas9 cassette and either a PMI or NPTII selectable marker.

Two plasmids were used for sorghum transformation to test the combination of *MiMe* editing and parthenogenesis (Supplemental Table 2). For *MiMe* editing tests, plasmid PHP103035 contained a Cas9 cassette, 4 gRNA cassettes (1 gRNA for each *MiMe* target gene, 2 gRNA driven by a maize U6 promoter and 2 gRNA driven by a sorghum U6 promoter) (Massel et al., 2022), and *NPTII* selectable marker. For a combined parthenogenesis and gene editing test, plasmid PHP105181 contained the maize *DD45:ASGR-BBML2* cassette, Cas9 cassette, 4 gRNA cassettes, and *NPTII* selectable marker.

### Sorghum transformation and transgenic edited event analyses

Immature embryo explants isolated from sorghum plants were transformed with *Agrobacterium* auxotrophic strain LBA4404 Thy-a carrying a ternary vector transformation system to generate transgenic sorghum plants (Anand et al., 2018; Che et al., 2018). Conventional, morphogenic gene-mediated, Wus2/CRE-mediated marker-free, and altruistic transformation methods were performed as previously described (Wu et al., 2014; Che et al., 2018; Che et al., 2022). The integrated copy number of the T-DNA and the vector backbone in these transgenic plants were determined by a series of qPCR analyses (Wu et al., 2014; Zhi et al., 2015). Amplicon deep sequencing to characterize CRISPR/Cas edits were performed as described (Che et al., 2022). Sequence reads were aligned to the Tx430, Tx623 or Macia wild-type reference sequence as appropriate via Bowtie2. The gRNA target sites, sequences, and corresponding PAM for *Sb-Spo111*, *Sb-Rec8*, *Sb-OsdL1,* and *Sb-OsdL3* genes were illustrated in Supplemental Figure 6. The primers used to amplify Sb-*Spo11*, Sb-*Rec8*, Sb-*OsdL1* and Sb-*OsdL3* genomic loci for amplicon analysis are shown in Supplemental Table 5.

### Cytological analyses

Sorghum reproductive tissues were collected before and after fertilization from sorghum plants. For analysis of parthenogenetic embryo development, fertilization was prevented by removing the stigmas from the immature ovules before anthesis and then ovules were collected 3 days after the stigma removal. A range of 15-25 ovaries from each plant were dissected using a LEICA MZ6 dissection microscope and then fixed in Carnoy’s fixative solution (3:1 ethanol:acetic acid) for 1 day at 4°C.

Ovaries were cleared in 100% methyl salicylate for at least 1 day. Cleared ovaries were observed under a LEICA DM5500 Q confocal microscope. Images were captured by LAS X software and further processed in Adobe Photoshop software. Male and female meiotic chromosome spreading was performed on meiotic stage floral buds fixed in Carnoy’s fixative solution for 1 day at 4°C and prepared as described previously (Ross et al., 1996). Chromosomes were stained with DAPI (4, 6-diamino-2-phenylindole dihydrochloride; 1.5 ug/ml) (Vector Laboratories, Inc. Burlingame, CA, USA) and observed with a LEICA DMRXA epifluorescence microscope system. Images were captured with LAS software and processed with Adobe Photoshop. Meiocytes were staged as described previously (Ross et al., 1996).

### Flow cytometry

Plant ploidy levels were measured using flow cytometry to detect the nuclear DNA content. Approximately 16mm^2^ fresh sorghum leaf tissue or pollen was added to a modified LB01 lysis buffer on ice (15 mM Tris, 2 mM Na_2_EDTA, 0.5 mM spermine 4HCl, 80 mM KCl, 20 mM NaCl, 0.1 % (v/v) Triton X-100, pH 7.5, 12.4 μM propidium iodide added before use, modified from prior studies (Dolezel & Bartos, 2005; Doležel et al., 1989), and was either chopped or lysed before being filtered through a 20μm filter. Nuclei suspensions were analyzed by the Attune^TM^ NxT Flow Cytometer and the corresponding Attune cytometric software (v5.3.0).

### SNP Marker analysis

SNP marker genotyping was performed using TaqMan SNP genotyping assays (ThermoFisher) using sorghum leaf samples as described previously (Ye et al., 2024).

## Funding

This work was supported in part by a sub-award from the Australian Commonwealth Scientific and Industrial Research Organisation (CSIRO) to Corteva under the Capturing Heterosis for Smallholder Farmers program (OPP1076280), and a sub-award from the University of Queensland to Corteva under the Hy-Gain program (INV-002955) from the Gates Foundation with supporting funds from Corteva Agriscience. The conclusions and opinions expressed in this work are those of the author(s) alone and shall not be attributed to the Foundation. Under the grant conditions of the Foundation, a Creative Commons Attribution 4.0 License has already been assigned to the Author Accepted Manuscript version that might arise from this submission. Please note works submitted as a preprint have not undergone a peer review process.

## Author Contributions

MKS, LY, PC, TJ, MCA designed the research. LY, KD and MKS analyzed the data. PC, LY, KD, MKS performed the experiments. MKS, MCA, IDG and AMGK wrote and revised the manuscript.

## Acknowledgements

We thank the members and advisors of the Capturing Heterosis and Hy-Gain collaborative teams. We additionally thank WeiWei Zhu and Amanda Reed for assistance with plant transformation, Sarah Anderson for assistance with manuscript preparation, and Corteva Agriscience Supervector, Genomics and Controlled Environments groups for support with sorghum materials, construct preparation and molecular analyses.

## Data Availability

Novel biological materials described in this publication may be available to the academic community and other not-for-profit institutions solely for non-commercial research purposes upon acceptance and signing of a material transfer agreement between the author’s institution and the requester. In some cases, such materials may originally contain genetic elements described in the manuscript that were obtained from a third party (for example, *DsRed*, *AmCyan*, and *Cas9*), and the authors may not be able to provide materials including third-party genetic elements to the requester because of certain third-party contractual restrictions placed on the author’s institution. In such cases, the requester will be required to obtain such materials directly from the third party. The authors and the authors’ institution do not grant any express or implied permission(s) to the requester to make, use, sell, offer for sale or import third-party proprietary materials. Obtaining any such permission(s) will be the sole responsibility of the requester. Corteva Agriscience proprietary germplasm will not be made available except at the discretion of Corteva Agriscience and then only in accordance with all applicable governmental regulations.

## Competing interests

The authors declare the following competing interests: MKS, LY, PC, KD, TJ, MCA are or have been employees of Corteva Agriscience.

## Supplemental Information

**Supplemental Figure 1.**
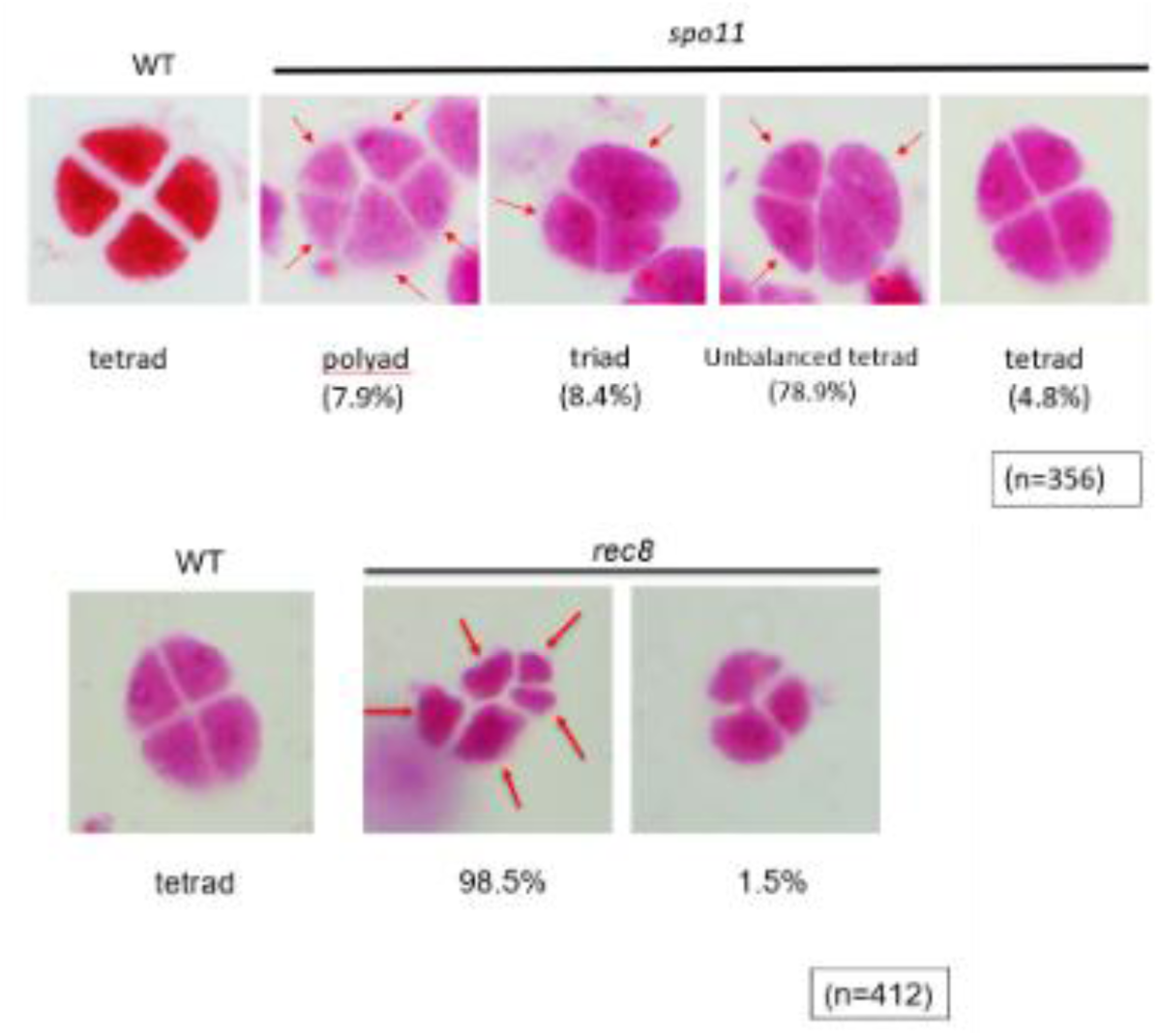
Tetrad analysis of *spo11-1* and *rec8* mutants in sorghum.

**Supplemental Figure 2.**
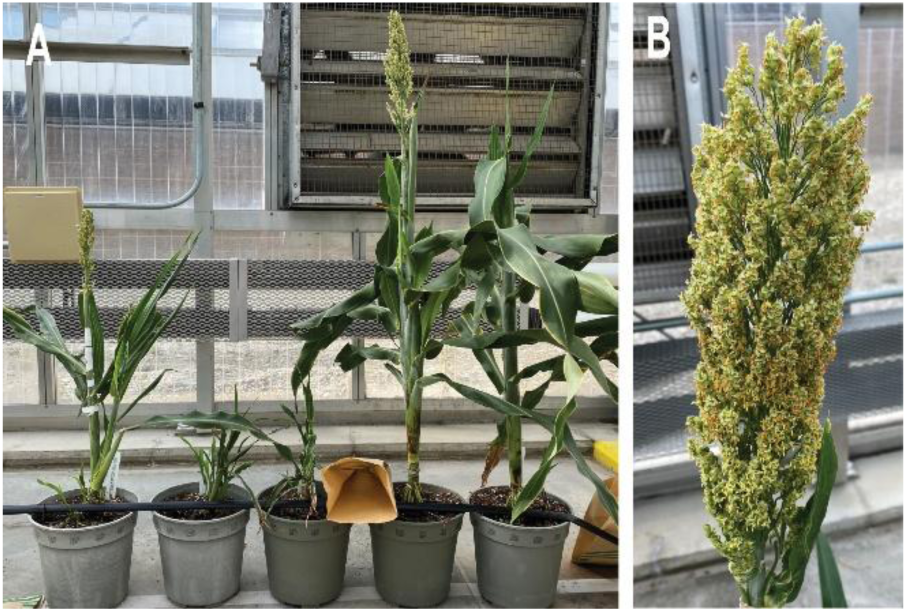
Two-step synthetic apomixis in Tx623/Tx430 hybrid sorghum. (A) Plant morphology of T0 events. (B) Panicle morphology of T0 event.

**Supplemental Figure 3.**
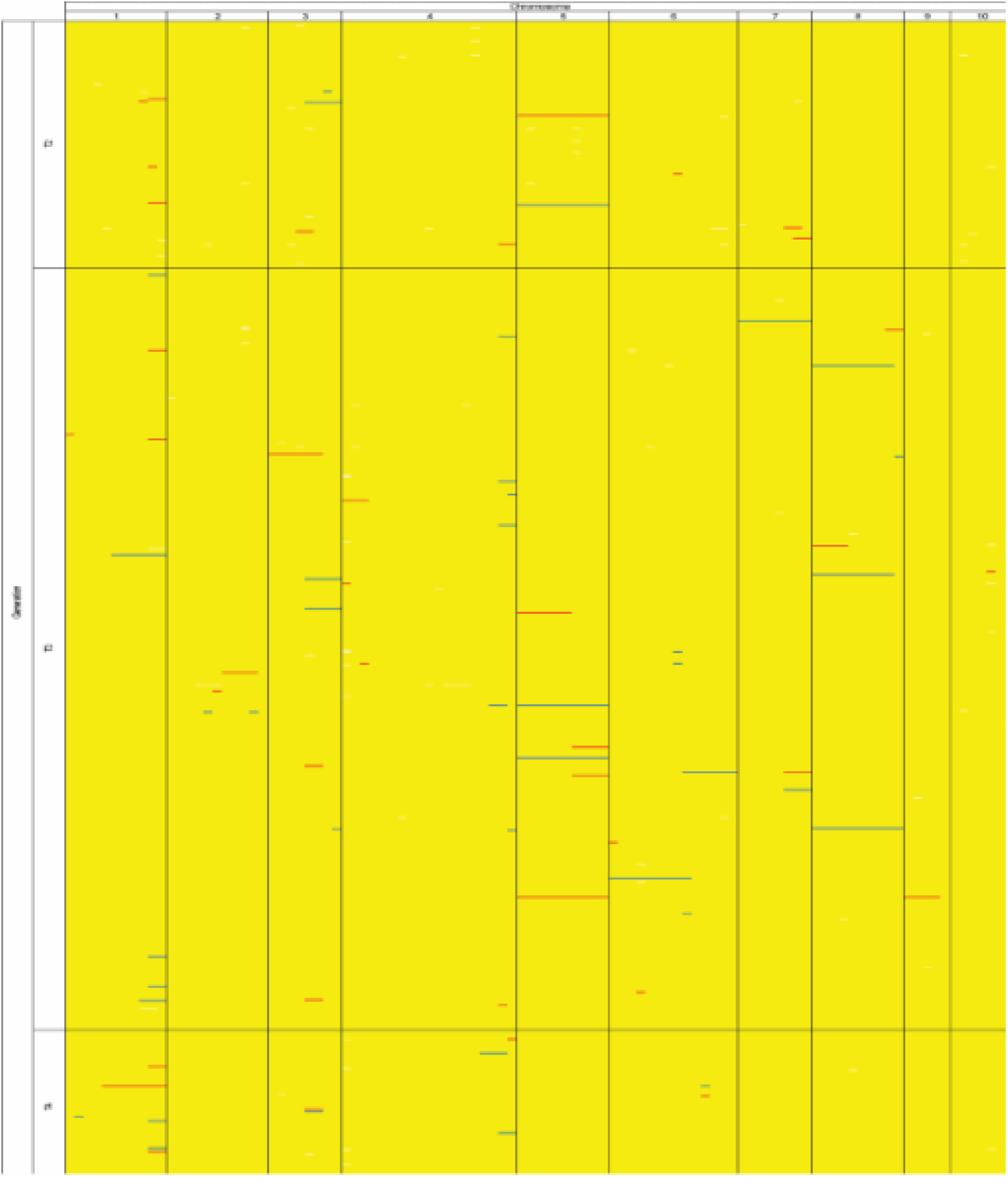
Multi-generation SNP marker analysis of selected event 143 of two-step synthetic apomixis in Tx623/Tx430 hybrid sorghum. The Tx430 alleles are marked in blue, Tx623 alleles in red, and heterozygous alleles in yellow

**Supplemental Figure 4.**
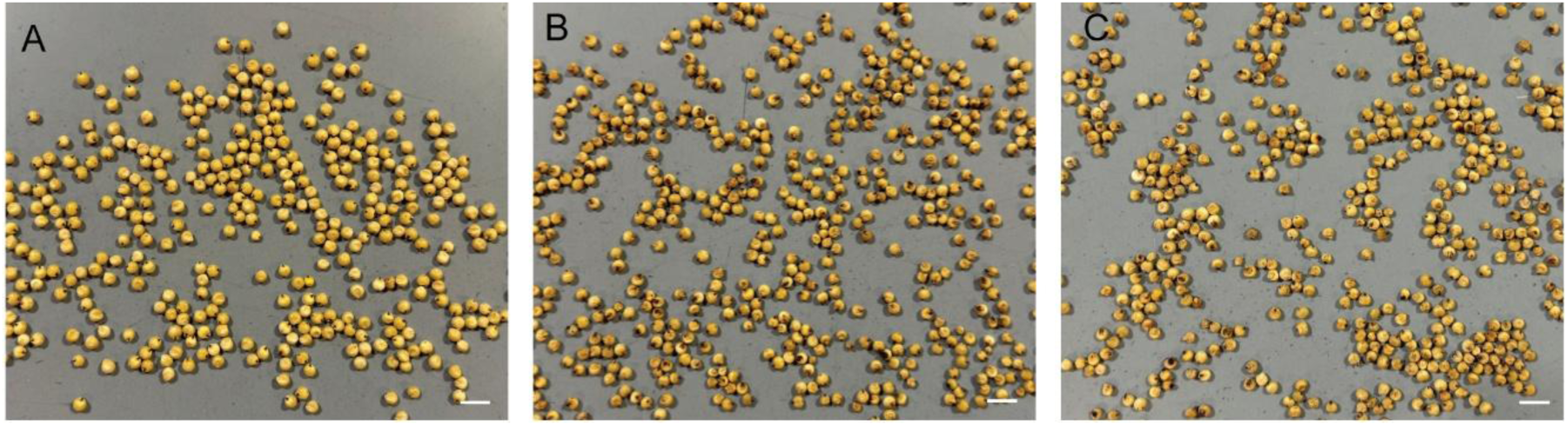
Representative seed of Tx623/Tx430 F1 hybrid control (A), T3 generation of event 143 apomictic hybrid (B), seed of T4 generation of event 143 apomictic hybrid (D), bar = 1cm.

**Supplemental Figure 5.**
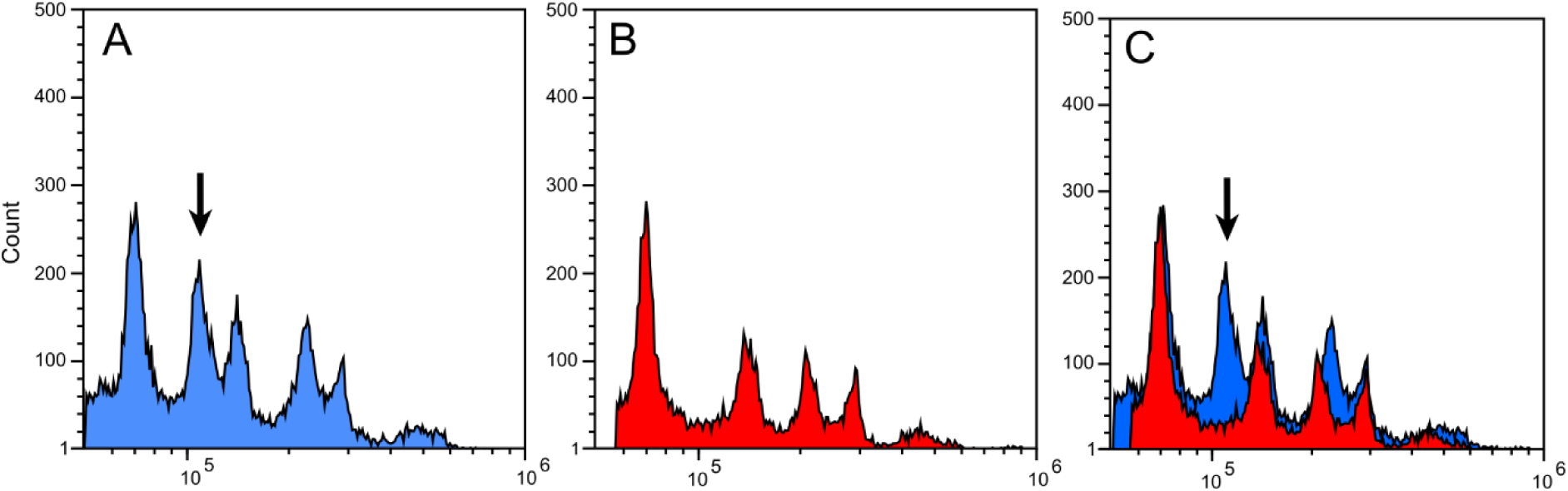
Representative flow cytometry histograms of PI-stained nuclei isolated from young developing kernels of WT control (A), Tx623/Tx430 apomictic event 143 plants (B) and an overlay of the two (C). WT control material contains diploid sporophytic tissue, a diploid immature embryo and triploid developing endosperm (arrow in A and C), whereas the apomictic material contains diploid sporophytic tissue, a diploid embryo and hexaploid developing endosperm. X axis is log relative fluorescent intensity (PI), Y axis is number of nuclei.

**Supplemental Figure 6.**
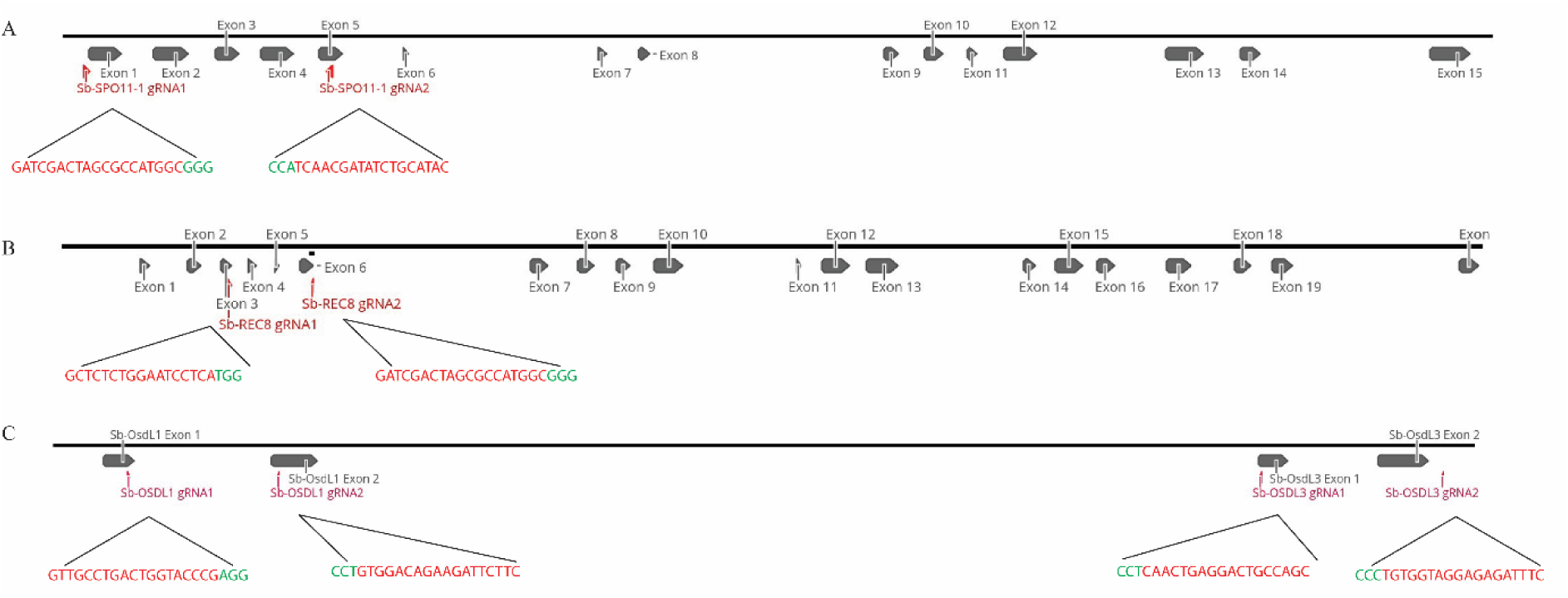
Diagram of *MiMe* loci gene structures and target sites. Gene structure of *Sb-Spo11-1* with gRNA target sites (in red) and PAM sequence (in green) (A), gene structure of *Sb-Rec8* with gRNA target sites (in red) and PAM sequence (in green) (B), gene structure of *Sb-OsdL1* and *Sb*-*OsdL3* with gRNA target sites (in red) and PAM sequence (in green) (C). *Geneious version 2025.0 created by Biomatters. Available from* https://www.geneious.com

**Supplemental Table 1.**
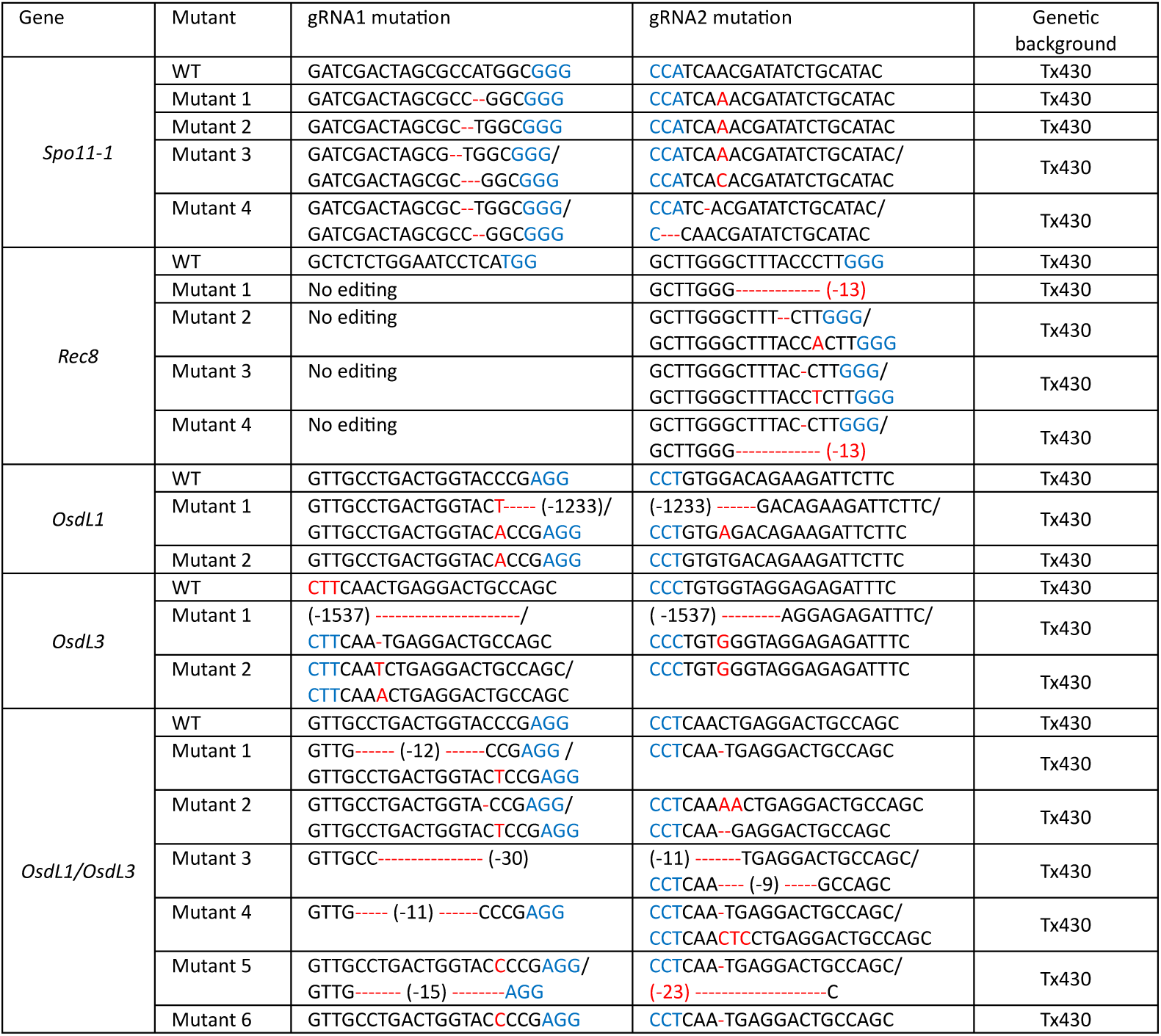
Alleles of *spo11-1*, *rec8*, *osdL1*, *osdL3*, and *osdL1 osdL3* in Tx430 background. PAM sequence indicated in blue, CRISPR/Cas9 induced insertion/deletion indicated in red.

**Supplemental Table 2.**
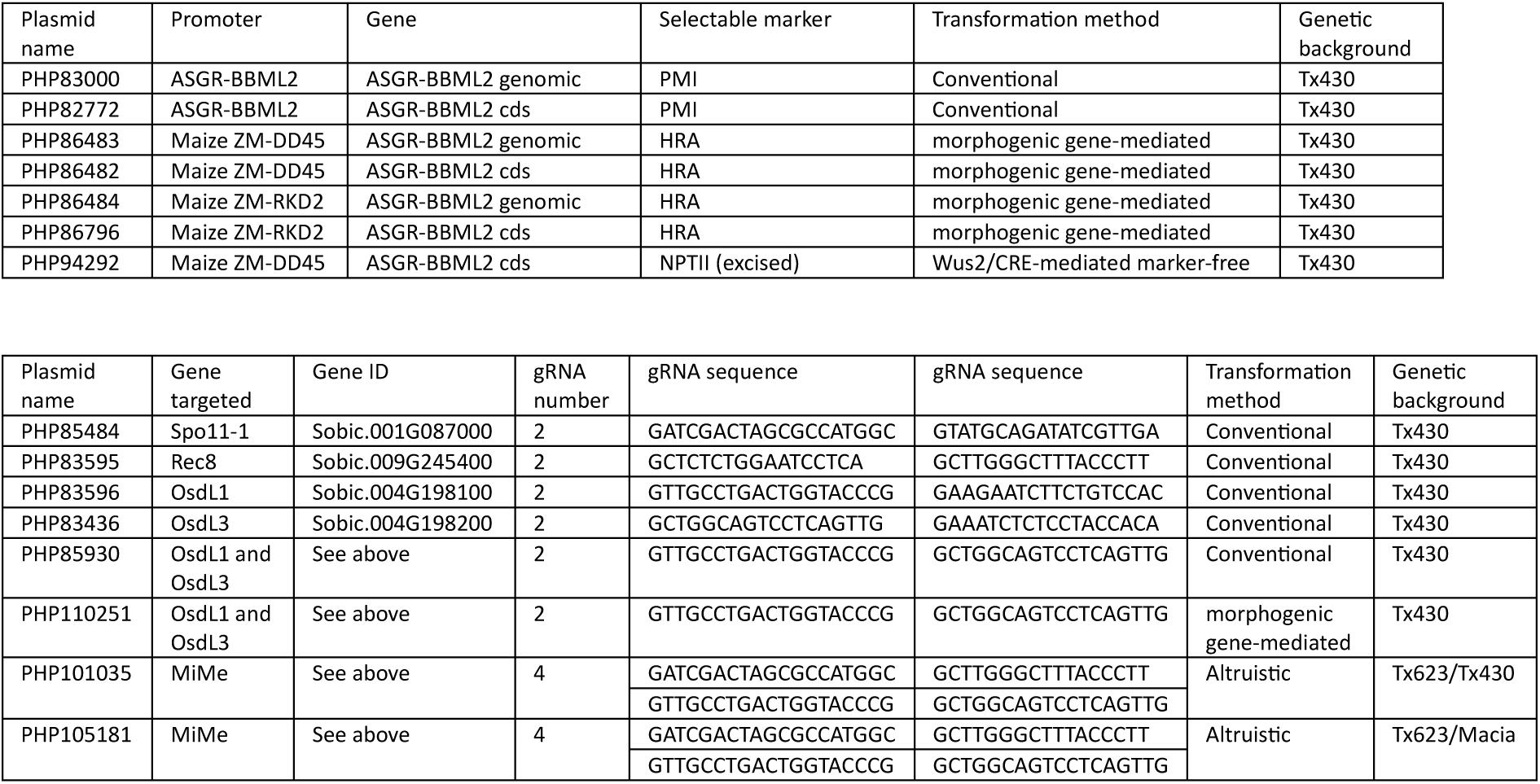
Plasmids used for transgenic and CRISPR/Cas9 editing, for Agrobacterium transformation

**Supplemental Table 3.** Characterization of Tx623/Tx430 and Tx623/Macia apomictic T0 CRISPR/Cas9 edit efficiencies at *OsdL1*, *OsdL3*, *Spo11-1* and *Rec8* target sites.

| Hybrid | T0 plants analyzed | Editing target site | Editing frequency | Biallelic frameshift allele editing frequency | Combined biallelic frameshift allele editing frequency |
| --- | --- | --- | --- | --- | --- |
| Tx623/Tx430 | 150 | OsdL1-CR3 | 45.3% | 24.7% | 13.3% |
|  |  | OsdL3-CR1 | 46.7% | 30.7% |  |
|  |  | Rec8-CR4 | 45.3% | 42% |  |
|  |  | Spo11-1-CR1 | 47.3% | 39.3% |  |
| Tx623/Macia | 160 | OsdL1-CR3 | 51.3% | 25.6% | 11.9% |
|  |  | OsdL3-CR1 | 53.1% | 28.1% |  |
|  |  | Rec8-CR4 | 51.9% | 45.6% |  |
|  |  | Spo11-1-CR1 | 51.9% | 48.8% |  |

**Supplemental Table 4.**
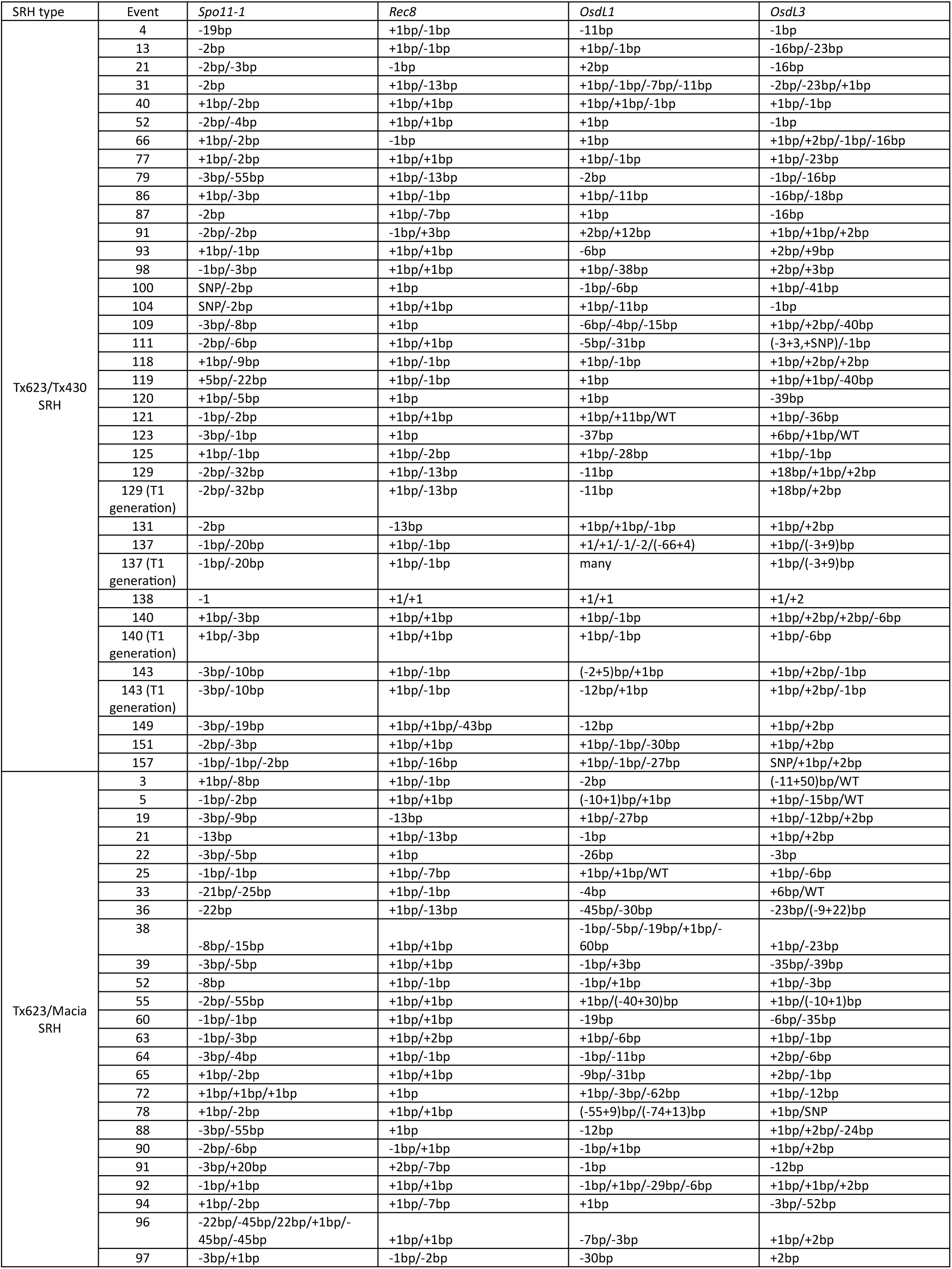
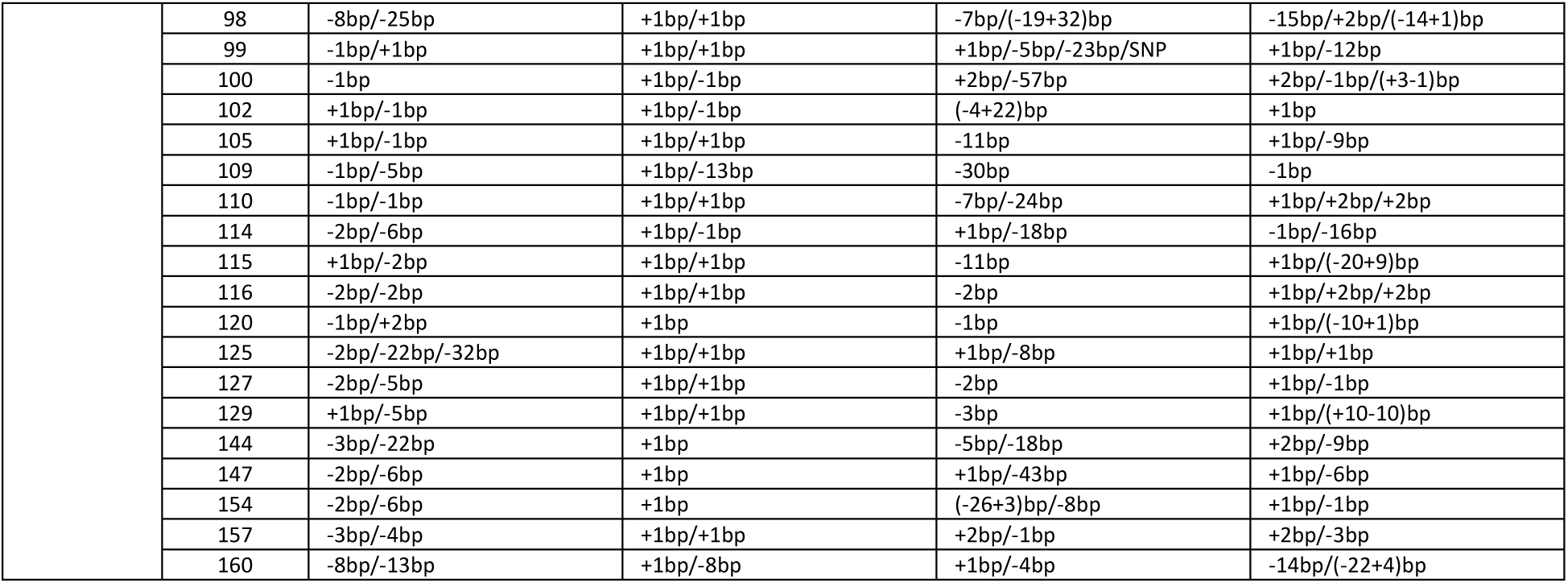
Characterization of Tx623/Tx430 apomictic T0 and T1 generation CRISPR/Cas9 edit alleles at *OsdL1*, *OsdL3*, *Spo11-1* and *Rec8* target sites. Analysis identified some T0 individuals that contained more than two unique alleles, suggesting chimerism within the T0 plant. * many unique alleles identified, each individual plant contained a single allele

**Supplemental Table 5.**
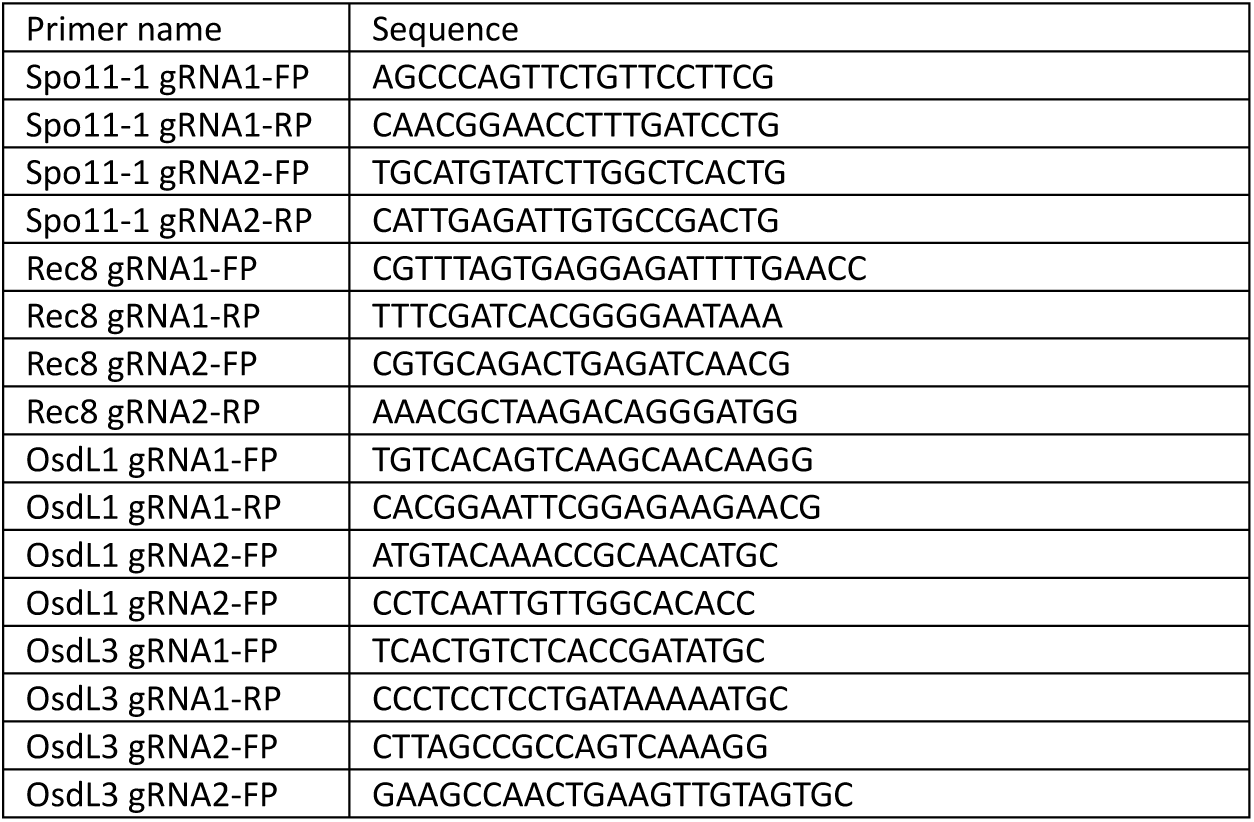
Primers sequences used for amplicon analysis

## References

Albertsen, M.C., Chamberlin, M.A.; Fox, T.W.; Lawit, S.J.; Loveland, B.R. (2013). Compositions and Methods for Expression of a Sequence in a Reproductive Tissue of a Plant. US20130180009A1

Anand, A., Bass, S.H., Wu, E., Wang, N., McBride, K.E., Annaluru, N., Miller, M., Hua, M., and Jones, T.J. (2018). An improved ternary vector system for Agrobacterium-mediated rapid maize transformation. Plant Mol Biol 97:187–200. 10.1007/s11103-018-0732-y.

Boutilier, K., Offringa, R., Sharma, V.K., Kieft, H., Ouellet, T., Zhang, L., Hattori, J., Liu, C.M., van Lammeren, A.A., Miki, B.L., et al. (2002). Ectopic expression of BABY BOOM triggers a conversion from vegetative to embryonic growth. Plant Cell 14:1737–1749. 10.1105/tpc.001941.

Che, P., Wu, E., Simon, M.K., Anand, A., Lowe, K., Gao, H., Sigmund, A.L., Yang, M., Albertsen, M.C., Gordon-Kamm, W., et al. (2022). Wuschel2 enables highly efficient CRISPR/Cas-targeted genome editing during rapid de novo shoot regeneration in sorghum. Communications Biology 5:344. 10.1038/s42003-022-03308-w.

Che, P., Anand, A., Wu, E., Sander, J.D., Simon, M.K., Zhu, W., Sigmund, A.L., Zastrow-Hayes, G., Miller, M., Liu, D., et al. (2018). Developing a flexible, high-efficiency Agrobacterium-mediated sorghum transformation system with broad application. Plant Biotechnol J 16:1388–1395. 10.1111/pbi.12879.

Conner, J.A., Podio, M., and Ozias-Akins, P. (2017). Haploid embryo production in rice and maize induced by PsASGR-BBML transgenes. Plant Reproduction 30:41–52. 10.1007/s00497-017-0298-x.

Conner, J.A., Mookkan, M., Huo, H., Chae, K., and Ozias-Akins, P. (2015). A parthenogenesis gene of apomict origin elicits embryo formation from unfertilized eggs in a sexual plant. Proceedings of the National Academy of Sciences 112:11205–11210. 10.1073/pnas.1505856112.

Dan, J., Xia, Y., Wang, Y., Zhan, Y., Tian, J., Tang, N., Deng, H., and Cao, M. (2024). One-line hybrid rice with high-efficiency synthetic apomixis and near-normal fertility. Plant Cell Reports 43:79. 10.1007/s00299-024-03154-6

de Wet, J.M.J. (1978). SPECIAL PAPER: SYSTEMATICS AND EVOLUTION OF SORGHUM SECT. SORGHUM (GRAMINEAE). American Journal of Botany 65:477–484. 10.1002/j.1537-2197.1978.tb06096.x.

d’Erfurth, I., Jolivet, S., Froger, N., Catrice, O., Novatchkova, M., and Mercier, R. (2009). Turning Meiosis into Mitosis. PLOS Biology 7:e1000124. 10.1371/journal.pbio.1000124.

Doležel, J., and Bartos, J. (2005). Plant DNA flow cytometry and estimation of nuclear genome size. Ann Bot 95:99–110. 10.1093/aob/mci005.

Doležel, J., Binarová, P., and Lcretti, S. (1989). Analysis of Nuclear DNA content in plant cells by Flow cytometry. Biologia Plantarum 31:113–120. 10.1007/BF02907241.

Fomicheva, M., Kulakov, Y., Alyokhina, K., & Domblides, E. (2024). Spontaneous and Chemically Induced Genome Doubling and Polyploidization in Vegetable Crops. Horticulturae, 10(6), 551. 10.3390/horticulturae10060551

Gilles, L.M., Khaled, A., Laffaire, J.B., Chaignon, S., Gendrot, G., Laplaige, J., Bergès, H., Beydon, G., Bayle, V., Barret, P., et al. (2017). Loss of pollen-specific phospholipase NOT LIKE DAD triggers gynogenesis in maize. The EMBO Journal 36:707–717. 10.15252/embj.201796603.

Hand, M.L., and Koltunow, A.M.G. (2014). The Genetic Control of Apomixis: Asexual Seed Formation. Genetics 197:441–450. 10.1534/genetics.114.163105.

Hanna, W.W., Bashaw, E.C. (1987). Apomixis: Its identification and use in plant breeding. Crop Science 27:1136–1139. 10.2135/cropsci1987.0011183X002700060010x

Heidemann, B., Primetis, E., Zahn, I.E., and Underwood, C.J. (2025). To infinity and beyond: recent progress, bottlenecks, and potential of clonal seeds by apomixis. The Plant Journal 121:e70054. 10.1111/tpj.70054.

Hojsgaard, D., Klatt, S., Baier, R., Carman, J.G., and Hörandl, E. (2014). Taxonomy and Biogeography of Apomixis in Angiosperms and Associated Biodiversity Characteristics. Critical Reviews in Plant Sciences 33:414–427. 10.1080/07352689.2014.898488.

Hossain, S., Islam, N., Rahman, M., Mostofa, M.G., and Khan, A.R. (2022). Sorghum: A prospective crop for climatic vulnerability, food and nutritional security. Journal of Agriculture and Food Research. 8:100300. 10.1016/j.jafr.2022.100300.

Huang, Y., Meng, X., Rao, Y., Xie, Y., Sun, T., Chen, W., Wei, X., Xiong, J., Yu, H., Li, J., et al. (2025). OsWUS-driven synthetic apomixis in hybrid rice. Plant Communications 6. 10.1016/j.xplc.2024.101136.

Iwata E, Ikeda S, Matsunaga S, Kurata M, Yoshioka Y, Criqui MC, Genschik P, Ito M. (2011). GIGAS CELL1, a novel negative regulator of the anaphase-promoting complex/cyclosome, is required for proper mitotic progression and cell fate determination in Arabidopsis. Plant Cell. 23(12):4382–93. doi: 10.1105/tpc.111.092049.

Iwata E, Ikeda S, Abe N, Kobayashi A, Kurata M, Matsunaga S, Yoshioka Y, Criqui MC, Genschik P, Ito M. (2012). Roles of GIG1 and UVI4 in genome duplication in Arabidopsis thaliana. Plant Signal Behav. 7(9):1079–81. 10.4161/psb.21133.

Kazungu, Florence Kadzo, Esther Mwende Muindi, and Jackson Muema Mulinge. (2023). “Overview of Sorghum (Sorghum Bicolor. L), Its Economic Importance, Ecological Requirements and Production Constraints in Kenya”. International Journal of Plant & Soil Science 35 (1):62–71. 10.9734/ijpss/2023/v35i12744.

Kelliher, T., Starr, D., Richbourg, L., Chintamanani, S., Delzer, B., Nuccio, M.L., Green, J., Chen, Z., McCuiston, J., Wang, W., et al. (2017). MATRILINEAL, a sperm-specific phospholipase, triggers maize haploid induction. Nature 542:105–109. 10.1038/nature20827.

Khalifa, M., and Eltahir, E.A.B. (2023). Assessment of global sorghum production, tolerance, and climate risk. Frontiers in Sustainable Food Systems 7. 10.3389/fsufs.2023.1184373.

Khanday, I., Skinner, D., Yang, B., Mercier, R., and Sundaresan, V. (2019). A male-expressed rice embryogenic trigger redirected for asexual propagation through seeds. Nature 565:91–95. 10.1038/s41586-018-0785-8.

Koltunow, A.M., and Grossniklaus, U. (2003). Apomixis: A Developmental Perspective. Annual Review of Plant Biology 54:547–574. 10.1146/annurev.arplant.54.110901.160842.

Leblanc, O., Grimanelli, D., Islam-Faridi, N., Berthaud, J., and Savidan, Y. (1996). Reproductive Behavior in Maize-Tripsacum Polyhaploid Plants: Implications for the Transfer of Apomixis Into Maize. Journal of Heredity 87:108–111. 10.1093/oxfordjournals.jhered.a022964.

Liu, C., He, Z., Zhang, Y., Hu, F., Li, M., Liu, Q., Huang, Y., Wang, J., Zhang, W., Wang, C., and Wang, K. (2023). Synthetic apomixis enables stable transgenerational transmission of heterotic phenotypes in hybrid rice. Plant Communications 4, 100470. 10.1016/j.xplc.2022.100470

Liu, C., Li, X., Meng, D., Zhong, Y., Chen, C., Dong, X., Xu, X., Chen, B., Li, W., and Li, L. (2017). A 4-bp insertion at ZmPLA1 encoding a putative phospholipase A generates haploid induction in maize. Molecular Plant 10:520–522. 10.1016/j.molp.2017.01.011

Marimuthu, M.P.A., Jolivet, S., Ravi, M., Pereira, L., Davda, J.N., Cromer, L., Wang, L., Nogué, F., Chan, S.W.L., Siddiqi, I., et al. (2011). Synthetic Clonal Reproduction Through Seeds. Science 331:876–876. 10.1126/science.1199682.

Massel, K., Lam, Y., Hintzsche, J., Lester, N., Botella, J.R., and Godwin, I.D. (2022). Endogenous U6 promoters improve CRISPR/Cas9 editing efficiencies in Sorghum bicolor and show potential for applications in other cereals. Plant Cell Reports 41:489–492. 10.1007/s00299-021-02816-z.

Mieulet, D., Jolivet, S., Rivard, M., Cromer, L., Vernet, A., Mayonove, P., Pereira, L., Droc, G., Courtois, B., Guiderdoni, E., et al. (2016). Turning rice meiosis into mitosis. Cell Res 26:1242–1254. 10.1038/cr.2016.117.

Nogler, G.A. (1984). Gametophytic Apomixis. In Embryology of Angiosperms, B.M. Johri, ed. (Springer Berlin Heidelberg: Berlin, Heidelberg), pp. 475–518. 10.1007/978-3-642-69302-1_10.

Ravi, M., and Chan, S.W. (2010). Haploid plants produced by centromere-mediated genome elimination. Nature 464:615–618. 10.1038/nature08842

Ren, H., Shankle, K., Cho, MJ. et al. (2024). Synergistic induction of fertilization-independent embryogenesis in rice egg cells by paternal-genome-expressed transcription factors. Nat. Plants 10, 1892–1899. 10.1038/s41477-024-01848-z

Richards, A.J. (2003). Apomixis in flowering plants: an overview. Philos Trans R Soc Lond B Biol Sci 358:1085–1093. 10.1098/rstb.2003.1294.

Ross KJ, Fransz P, Jones GH. (1996). A light microscopic atlas of meiosis in Arabidopsis thaliana. Chromosome Res. Nov;4(7):507–16. doi: 10.1007/BF02261778.

Song, M., Li, F., Chen, Z., Hou, H., Wang, Y., Liu, H., Liu, D., Li, J., Peng, T., Zhao, Y., et al. (2024). Engineering high-frequency apomixis with normal seed production in hybrid rice. iScience 2710.1016/j.isci.2024.111479.

Song M, Wang W, Ji C, Li S, Liu W, Hu X, Feng A, Ruan S, Du S, Wang H, Dai K, Guo L, Qian Q, Si H, Hu X. (2024). Simultaneous production of high-frequency synthetic apomixis with high fertility and improved agronomic traits in hybrid rice. Molecular Plant 17(1):4–7. 10.1016/j.molp.2023.11.007.

Underwood, C.J., and Mercier, R. (2022). Engineering Apomixis: Clonal Seeds Approaching the Fields. Annual Review of Plant Biology 73:201–225. 10.1146/annurev-arplant-102720-013958.

Underwood, C.J., Vijverberg, K., Rigola, D., Okamoto, S., Oplaat, C., Camp, R.H.M.O.d., Radoeva, T., Schauer, S.E., Fierens, J., Jansen, K., et al. (2022). A PARTHENOGENESIS allele from apomictic dandelion can induce egg cell division without fertilization in lettuce. Nature Genetics 54:84–93. 10.1038/s41588-021-00984-y.

Vernet, A., Meynard, D., Lian, Q., Mieulet, D., Gibert, O., Bissah, M., Rivallan, R., Autran, D., Leblanc, O., Meunier, A.C., et al. (2022). High-frequency synthetic apomixis in hybrid rice. Nature Communications 13:7963. 10.1038/s41467-022-35679-3.

Wang, Y., Fuentes, R.R., van Rengs, W.M., Effgen, S., Zaidan, M.W.A.M., Franzen, R., Susanto, T., Fernandes, J.B., Mercier, R., and Underwood, C.J. (2024). Harnessing clonal gametes in hybrid crops to engineer polyploid genomes. Nature Genetics 56:1075–1079. 10.1038/s41588-024-01750-6

Wang, C., Liu, Q., Shen, Y., Hua, Y., Wang, J., Lin, J., Wu, M., Sun, T., Cheng, Z., Mercier, R., et al. (2019). Clonal seeds from hybrid rice by simultaneous genome engineering of meiosis and fertilization genes. Nature Biotechnology 37:283–286. 10.1038/s41587-018-0003-0.

Wei, X., Liu, C., Chen, X., Lu, H., Wang, J., Yang, S., and Wang, K. (2023). Synthetic apomixis with normal hybrid rice seed production. Molecular Plant 16:489–492. 10.1016/j.molp.2023.01.005

Wu, E., Lenderts, B., Glassman, K., Berezowska-Kaniewska, M., Christensen, H., Asmus, T., Zhen, S., Chu, U., Cho, M.J., and Zhao, Z.Y. (2014). Optimized Agrobacterium-mediated sorghum transformation protocol and molecular data of transgenic sorghum plants. In Vitro Cell Dev Biol Plant 50:9–18. 10.1007/s11627-013-9583-z.

Xie, E., Li, Y., Tang, D., Lv, Y., Shen, Y., and Cheng, Z. (2019). A strategy for generating rice apomixis by gene editing. Journal of Integrative Plant Biology 61:911–916. 10.1111/jipb.12785.

Ye, H., Louden, M., and Reinders, J.A.T. (2024). A novel in vivo genome editing doubled haploid system for Zea mays L. Nature Plants 10:1493–1501. 10.1038/s41477-024-01795-9.

Zhi, L., TeRonde, S., Meyer, S., Arling, M.L., Register Iii, J.C., Zhao, Z.-Y., Jones, T.J., and Anand, A. (2015). Effect of Agrobacterium strain and plasmid copy number on transformation frequency, event quality and usable event quality in an elite maize cultivar. Plant Cell Reports 34:745–754. 10.1007/s00299-014-1734-0.

